# Gamma oscillations in basal ganglia stem from the interplay between local inhibition and beta synchronization

**DOI:** 10.1101/2025.03.19.644161

**Authors:** Federico Fattorini, Mahboubeh Ahmadipour, Enrico Cataldo, Alberto Mazzoni, Nicolò Meneghetti

## Abstract

Basal ganglia rhythms have mainly been studied in the beta band (12–30 Hz), a hallmark of Parkinson’s disease (PD), while gamma oscillations (30–100 Hz) in the subthalamic nucleus (STN) have emerged as alternative markers for guiding adaptive deep brain stimulation. However, their underlying mechanisms remains unclear. Using a spiking network model of the basal ganglia, we identified two distinct gamma rhythms: a high-frequency gamma in pallidal (GPe-TI) neurons and a slower gamma in D2 medium spiny neurons (MSNs), both generated through self-inhibition. Under simulated parkinsonian condition, GPe-TI gamma intensity remained stable. In contrast, D2 MSN gamma emerged only in pathological conditions and was strongly modulated by beta activity in both intensity and frequency. Although STN did not generate gamma oscillations directly, gamma activity from GPe-TI population was reflected in simulated STN local field potentials. These results clarify the circuit origins of gamma rhythms and their modulation in PD.

## Introduction

The basal ganglia (BG) are a group of interconnected subcortical nuclei in the brain that play a pivotal role in regulating motor control, learning, emotions, and decision-making. The main structures that comprise the basal ganglia are the striatum (STR), globus pallidus (GP), subthalamic nucleus (STN), and substantia nigra (SN) ^1^. The striatum contains fast-spiking interneurons (FSNs) and medium-spiny neurons (MSNs), which express D1- and D2-type dopamine receptors, while the GP is subdivided into the external (GPe) and internal (GPi) segments. These nuclei are interconnected in a complex network that helps modulate voluntary movement and other cognitive functions ^2^. Disruptions in the functioning of these structures, particularly in the context of neurodegenerative diseases, can lead to profound motor and behavioral impairments. Parkinson’s disease (PD) is the second most common neurodegenerative disorder, affecting approximately 0.3% of the global population. The disease is primarily characterized by motor symptoms such as akinesia ^3^, bradykinesia ^4^, resting tremor ^5^, and gait disturbances ^6^. PD is caused by the progressive degeneration of dopaminergic neurons in the substantia nigra pars compacta ^7^, leading to a depletion of dopamine in the striatum and disruptions in basal ganglia signaling. This depletion specifically affects D1- and D2-receptor-expressing neurons, which alters the dynamic balance of activity within the basal ganglia network. As PD progresses, these disruptions manifest as motor impairments and are often managed through pharmacological treatments or surgical interventions. One such intervention is deep brain stimulation (DBS), a surgical approach that delivers electrical pulses to specific basal ganglia nuclei, typically the STN or GPi, to alleviate motor symptoms ^8,9^. However, conventional DBS delivers continuous stimulation without accounting for fluctuations in symptom severity or variations in the patient’s daily activities. This limitation has spurred the development of adaptive DBS (aDBS), a more advanced form of stimulation that adjusts in real-time based on the patient’s physiological signals ^10–12^. aDBS requires continuous monitoring of brain activity to modulate stimulation parameters in response to dynamic changes in motor function. A critical challenge for aDBS is identifying electrophysiological biomarkers that reliably correlate with symptoms, reflect pathophysiological mechanisms, and support precise real-time modulation of electrical stimulation.

Among the electrophysiological changes observed in PD ^13–17^, oscillatory activity in the basal ganglia has been the subject of intense study. Of these oscillations, beta-band activity (12–30 Hz) has garnered the most attention. Beta oscillations are prominently observed in the STN and GPi of PD patients, and their amplitude correlates with the severity of motor impairment ^18–21^. Moreover, beta oscillations were found in both rats ^22^ and monkeys ^23,24^ models of PD, confirming the key role of these oscillations in the disease. These oscillations are now widely considered as biomarkers for DBS, particularly for adjusting aDBS ^11^. However, beta-band oscillations are influenced by various factors, including DBS stimulation, medication, and task performance, which can complicate their use in real-time modulation ^25–27^. This inherent variability has led to growing interest in gamma oscillations (30–100 Hz) as an alternative, more stable biomarker for adaptive DBS ^28–30^.

Gamma oscillations have emerged as a significant biomarker across a range of pathological conditions ^31–33^, including PD. These oscillations have been observed in the STN and other basal ganglia structures in both PD patients ^34–41^ and animal models ^42–45^. They are modulated by movement ^46,47^, associated with dyskinetic states ^48^, and linked to sleep disturbances commonly observed in PD ^49^. Unlike beta oscillations, gamma rhythms show a stronger robustness to external influences such as DBS stimulation and medication, making them a promising candidate for real-time monitoring in aDBS systems ^29,30^. Moreover, recent studies have demonstrated that gamma oscillations are tightly correlated with motor dysfunction ^35^, suggesting their potential role in PD symptomatology.

Despite the growing interest in gamma oscillations, the precise mechanisms underlying their generation and modulation in the basal ganglia network are not fully understood. Computational modeling, which has been used extensively to study basal ganglia dynamics, serves as a valuable tool to address these open problems ^50–52^. While some computational models have explored gamma rhythms in the BG, most have focused on beta oscillations and treated gamma as a secondary epiphenomenon. For example, Hodgkin-Huxley-based models have shown that gamma oscillations (∼80 Hz) can emerge in FSNs and GPe, with STN-DBS restoring these rhythms following dopamine depletion ^53^. Other studies have highlighted specific BG loops, such as the GPe-STN network, as key players in generating gamma rhythms in the 35–50 Hz range ^54,55^. Additionally, spiking models have linked gamma activity in the STN and GPe to motor control tasks ^56^ or variations in MSN connectivity ^57^. However, the role of gamma oscillations within the broader BG circuitry remains poorly understood.

In this study, we extend a previously established computational model ^58^ to explore the dynamics of gamma oscillations in the basal ganglia. We focus on the role of local inhibition in driving gamma rhythms in the striatum and GPe and examine how dopamine depletion, synaptic parameters, and external input strengths influence these oscillations. Our work aims to provide new insights into the mechanisms of gamma oscillations in PD and propose a theoretical framework for their potential use as biomarkers in adaptive DBS systems.

## Results

### Gamma oscillations in a computational model of basal ganglia

We investigated dopamine-depletion-driven oscillations in the basal ganglia (BG) firing activity starting from a previously validated spiking network model ^58^ (see Methods and Figure 1A). Coherently with previous findings ^58^, beta-range oscillations (12–30 Hz) were well-defined across all BG nuclei under healthy conditions (dopamine depletion level, D_d_ = 0.75; black lines in Figure 1B). Dopamine depletion enhanced the power of these beta oscillations in all nuclei (D_d_ = 1; orange lines in Figure 1B and Figure S2A), reproducing the emergence of pathological beta activity observed in the PD condition.

**Figure 1.**
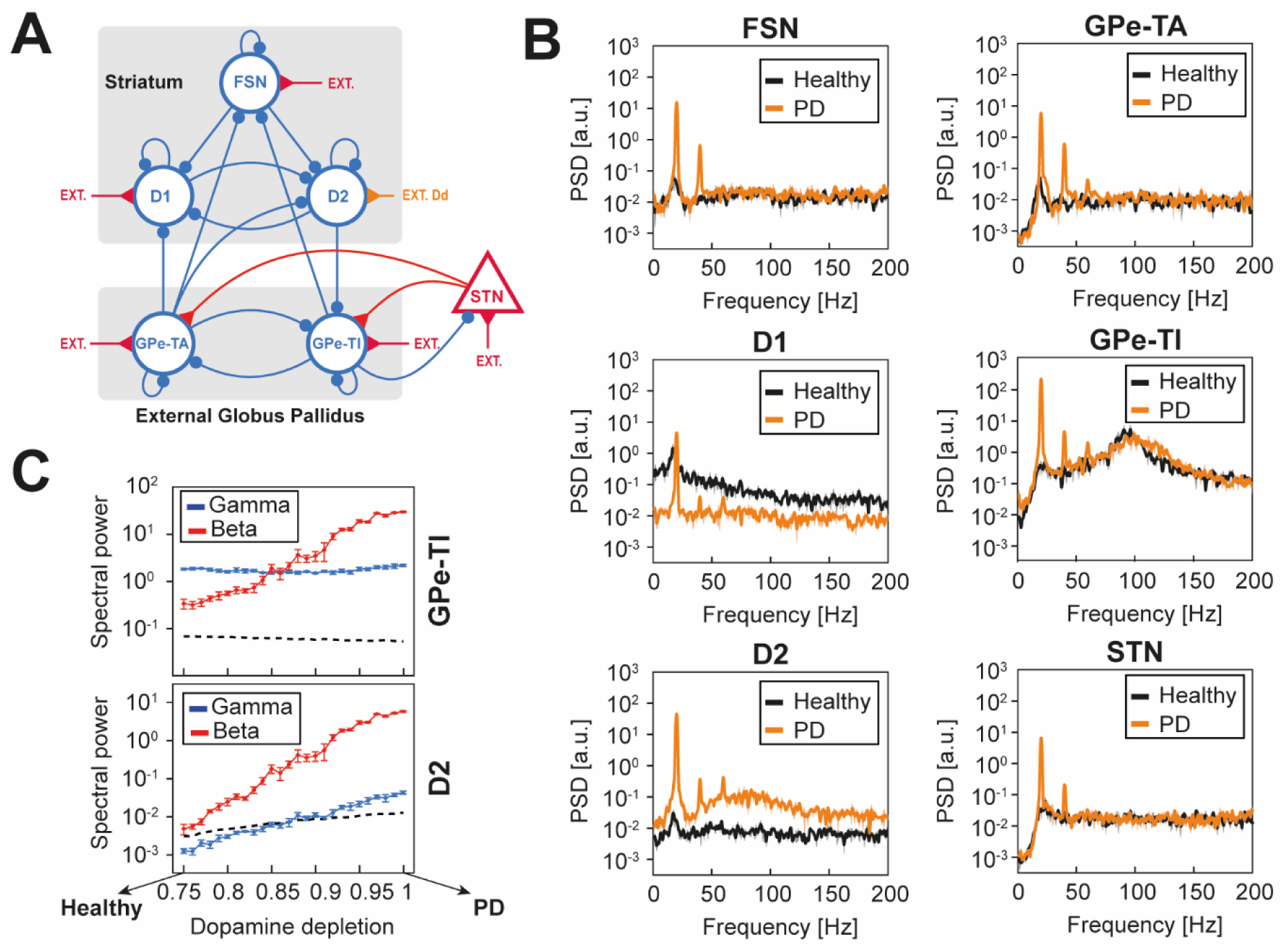
Gamma oscillations in the globus pallidus and the striatum of a basal ganglia spiking model. (A) Architecture of the BG model. FSN, D1 and D2: striatal spiking interneurons, and medium spiny neurons with D1 and D2 dopamine receptors; GPe-TA and GPe-TI: arkypallidal and prototypic populations of the globus pallidus pars externa; STN: subthalamic nucleus; ext: external excitatory Poissonian input. Red/blue arrows indicate excitatory/inhibitory projections. Dopamine depletion (D_d_, orange external drive) was modeled by adjusting the excitatory Poisson input to D2 neurons. (B) Power spectral densities (PSDs) of each BG nucleus activity under healthy (D_d_= 0.75, black) and Parkinson’s disease conditions (D_d_= 1, orange). PSDs are reported as mean and standard deviation across five simulations of 5 seconds. (C) Spectral powers of GPe-TI (top) and D2 (bottom) firing activity within beta (red) and gamma (blue) ranges across dopamine depletion levels. Data are reported as mean and standard deviation across five simulations. The threshold for statistically significant oscillatory power (dashed black line) was determined considering the power spectrum of an equivalent Poisson process with matched firing rate.

In contrast to the widespread beta oscillations, activity in the high gamma range (∼100 Hz) was observed only in the GPe-TI nucleus under healthy conditions (Figure 1B). Under dopamine-depleted states, gamma oscillations persisted in GPe-TI and newly emerged in the D2 population (Figure 1B). Interestingly, gamma oscillations in GPe-TI showed little to no change in power with increasing dopamine depletion (Figure 1C; Spearman ρ = 0.19, p = 0.36), while their peak frequency exhibited a non-monotonic trend (Figure S2B, bottom). In contrast, gamma activity in the D2 population was highly sensitive to dopamine depletion: gamma power increased significantly with greater depletion levels (Figure 1C; Spearman ρ = 0.99, **p << 0.001), with significant oscillations emerging only for D_d_ > 0.87 and showing an increase in peak frequency at higher depletion levels (Figure S2C, bottom).

These findings highlight two key insights. First, gamma oscillations in the D2 population are sensitive to dopamine dynamics, suggesting they may reflect pathological changes associated with Parkinson’s disease progression. Second, GPe-TI and D2 populations exhibit differential sensitivity to dopamine depletion, with gamma oscillations in GPe-TI being more resilient, while those in D2 increase in intensity and frequency with dopamine deficiency. Furthermore, our results reveal a distinct relationship between beta- and gamma-range oscillatory dynamics during dopamine depletion. In the following sections, we will explore the mechanistic origins of these oscillations and discuss their implications for understanding the emergence and regulation of abnormal rhythmic activity in the basal ganglia during PD progression.

### Globus pallidus self-inhibition drives gamma oscillations features

To explore the role of specific synaptic pathways in generating GPe-TI gamma oscillations within our basal ganglia model, we systematically silenced each connection individually and analyzed the resulting effects on GPe-TI gamma power (see Methods). Specifically, we calculated the ratio of GPe-TI gamma power with each connection silenced to the GPe-TI gamma power in the intact full-BG model (Figure 2A). Our results demonstrated that gamma oscillations were still present after the removal of most connections (Figure 2A), with one notable exception: GPe-TI self-inhibition. Disrupting this self-inhibitory pathway led to a near-complete loss of gamma oscillatory activity (Figure 2B), underscoring GPe-TI self-inhibition as a necessary mechanism for gamma rhythmogenesis within this nucleus. Interestingly, the removal of GPe-TI self-inhibition was also associated with a marked increase in beta oscillations power (Figure 2B).

**Figure 2.**
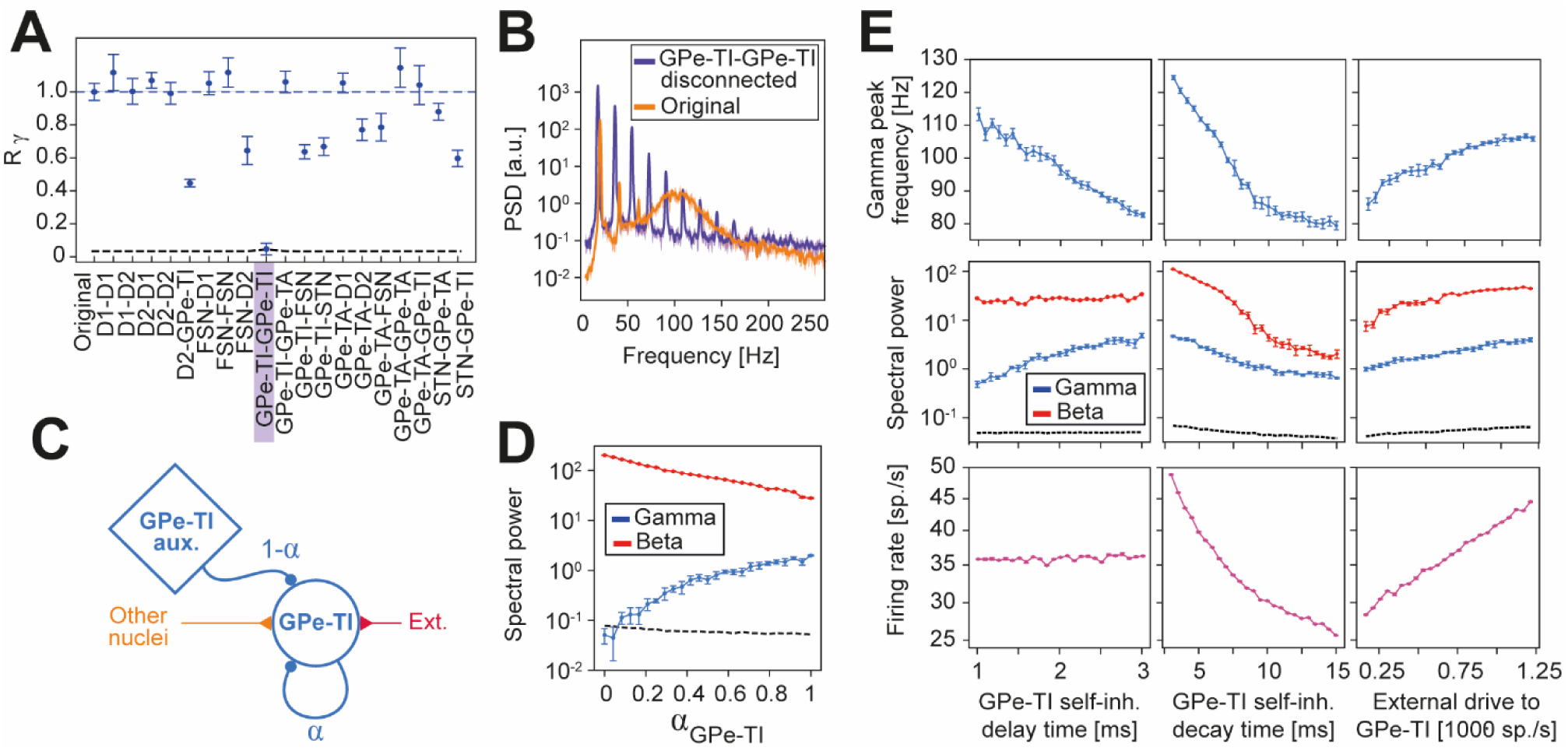
Self-inhibition parameters and external drive modulate GPe-TI oscillations. (A) Ratio R_ɣ_ of GPe-TI gamma power with each synaptic connection silenced compared to the GPe-TI gamma power in the intact full-BG model. The spectral power ratio of GPe-TI activity in the gamma band is shown with (‘original’) and without each synaptic connection in the full-BG model, with the specific connection removed indicated on the x-axis. For each removed connection, the threshold (black dashed line) for significant oscillations is computed as the ratio between the power of equivalent poisson processes and the gamma power in the fully connected model. The blue dashed line indicates the spectral power ratio for the fully connected case. (B) Power spectral densities (PSDs) of GPe-TI with (orange) and without (purple) self-inhibitory connection, in Parkinsonian condition (D_d_=1). (C) Scheme illustrating the procedure for reducing GPe-TI self-inhibition: the weights of self-inhibitory connections were progressively decreased by a multiplicative factor α Є [0,1], and input from an auxiliary Poissonian population (GPe-TI aux.) was added to compensate for the reduction in self-inhibitory input. The afferent activity from other BG nuclei and external inputs were left unchanged. (D) GPe-TI spectral power of beta (red) and gamma (blue) bands across self-inhibition synaptic strength α. The threshold for statistically significant oscillatory power (dashed black line) was determined considering the power spectrum of an equivalent Poisson process with matched firing rate. (E) Gamma peak frequency (top row), beta (red) and gamma (blue) spectral powers (middle row), and GPe-TI firing rate (bottom row) across GPe-TI self-inhibition delay (left column), GPe-TI self-inhibition decay time (middle column), and the strength of external input to GPe-TI (right column). In the middle row, the threshold for significant oscillations was indicated with a black dashed line. All data was reported as mean and standard deviation across five simulations.

To further investigate the role of GPe-TI self-inhibition in regulating gamma oscillations, we systematically reduced the strength of self-inhibitory feedback and assessed its effect on GPe-TI gamma power. To maintain a consistent average input to GPe-TI despite the reduced self-inhibition, we introduced an auxiliary Poisson inhibitory input (see Methods and Figure 2C, where it is referred to as "GPe-TI aux."). However, this compensatory input was insufficient to fully stabilize the average firing rate of the GPe-TI population, which consequently increased as the strength of self-inhibition decreased (Figure S3A, left).

Our findings revealed that progressively weakening self-inhibition synaptic strength α_Gpe-TI_, led to a gradual reduction in gamma activity (Figure 2D, Spearman ρ = 1.00, **p<0.001) and peak frequency (Figure S3A, right and Spearman ρ = 0.86, **p<0.001). When self-inhibition was fully silenced, gamma activity disappeared, declining to levels of power below statistical significance (Figure 2D). Notably, we also observed a concurrent increase in beta synchronization as self-inhibition strength decreased (Figure 2D, Spearman ρ = −0.96, **p<0.001) suggesting an inverse relationship between GPe-TI gamma oscillations (or, equivalently, self-inhibition strength) and beta hypersynchrony.

We next examined how other characteristics of GPe-TI self-inhibition (specifically, synaptic decay time and delay) impacted gamma activity. We found that the increase of both delay and decay times of GPe-TI self-inhibition synapses reduced the gamma peak frequency (Figure 2E top-left and top-middle. Delay: Spearman = −0.99, **p<0.001. Decay time: Spearman ρ = −0.99, **p<0.001), though their effects on gamma power differed: increasing synaptic delay boosted gamma power (Figure 2E middle-left, Spearman ρ = 0.98, **p<0.001), whereas increasing decay time reduced it (Figure 2E middle, Spearman ρ = −0.98, **p<0.001). Notably, the reduction in gamma power associated with increased decay time was accompanied by a decrease in the average firing rate of GPe-TI (Figure 2E bottom-middle; Spearman ρ = −1.00, **p<0.001). In contrast, the increase in gamma power due to prolonged synaptic delays occurred without a corresponding increase in GPe-TI firing rate (Figure 2E bottom-left, Spearman ρ = 0.54, *p = 0.005), suggesting that this enhancement reflected improved synaptic efficacy rather than increased firing activity alone.

We also observed distinct effects of decay time and delay on beta oscillations. Interestingly, we found beta oscillations power to closely reflect the average GPe-TI firing activity. GPe-TI self-inhibition delay had little to no effect on both beta power (Figure 2E middle-left, Spearman ρ = 0.53, *p = 0.007) and firing rate. Conversely, increased decay times significantly reduced both beta and GPe-TI firing (Figure 2E middle, Beta power: Spearman = −0.99, **p<0.001). Taken together, these findings suggest that while beta oscillations follow the average firing rate of GPe-TI, gamma oscillations are influenced not only by firing rate but also by local resonance properties within GPe-TI that are not reflected in beta oscillations.

Finally, we examined the effect of excitatory drive on GPe-TI gamma activity. As expected, increasing the excitatory drive to GPe-TI raised its average firing rate (Figure 2E bottom-right Spearman ρ = 1.00, **p<0.001), which in turn elevated both beta and gamma power (Figure 2E middle-right. Beta power: Spearman = 0.99, **p<0.001. Gamma power: Spearman ρ = 0.99, **p<0.001), as well as the gamma peak frequency (Figure 2E top-right, Spearman ρ = 0.99, **p<0.001). This finding underscores a link between GPe-TI firing activity and gamma oscillatory dynamics, with excitatory input modulating gamma features primarily through its effect on firing rate. However, as shown earlier, gamma oscillations also depend on intrinsic local dynamics within GPe-TI, suggesting that gamma power reflects a combination of both firing rate and the synaptic properties of the recurrent inhibition.

### Gamma oscillations in the globus pallidus are modulated by beta activity

In the previous section, we demonstrated that GPe-TI self-inhibition was necessary for the presence of gamma oscillations in this nucleus. Here, we assessed whether this self-inhibitory loop alone was also sufficient for gamma rhythmogenesis. To this end, we analyzed an isolated GPe-TI population, configured with the same parameters as the original network but disconnected from all other basal ganglia nuclei. Accordingly, the inputs from other nuclei were replaced with auxiliary constant-rate Poissonian inputs that matched the firing activity of their respective original projections (in Figure 3A, indicated with “GPe-TA aux.”, “D2 aux.” and “STN aux.”).

**Figure 3.**
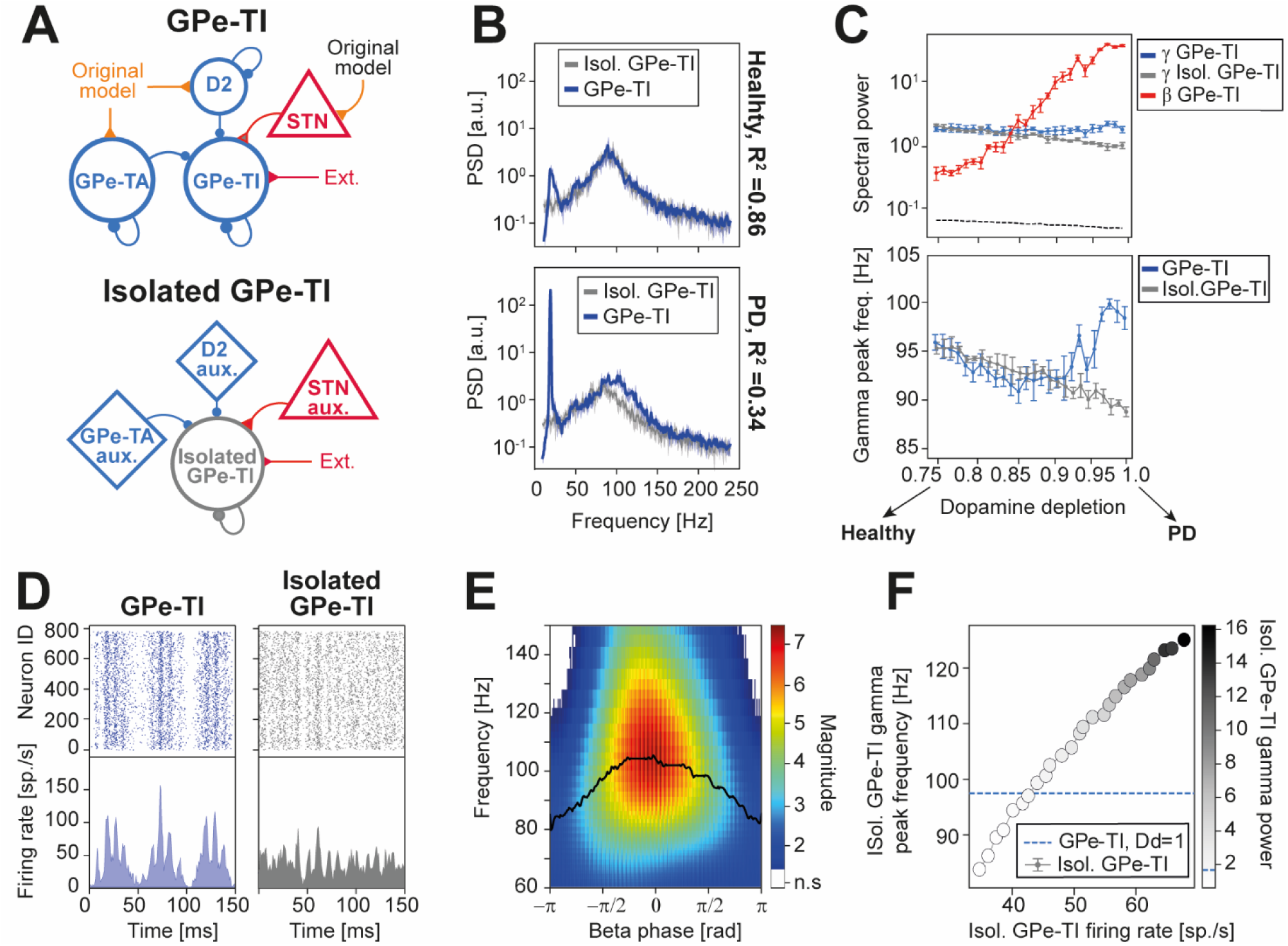
Interplay between beta and gamma oscillations in GPe-TI. (A) Schematic of isolation procedure for GPe-TI: the original nucleus (top) was detached by the rest of the model and the inputs from other nuclei were substituted with Poissonian sources (indicated with “aux.” in the bottom panel) with the same firing rates. External drive and self-inhibition were left unchanged. (B) Comparison of the power spectral densities (PSDs) of GPe-TI (blue) and isolated GPe-TI (grey) under healthy (D_d_=0.75, bottom) and Parkinsonian conditions (D_d_=1, bottom). (C) Spectral power (top) and gamma peak frequency (bottom) of GPe-TI (blue) and isolated GPe-TI (grey) across D_d_. The spectral power in the beta band (red) and the threshold for significant oscillations (black dashed lines), computed as the spectral power of a Poissonian process with the same firing rate, are also reported. Results are reported as mean across five simulations of 5 seconds. (D) Raster plots (top) and activity traces (bottom) of GPe-TI (left) and isolated GPe-TI (right), in Parkinsonian condition. (E) Scalogram of GPe-TI activity averaged across beta cycles, under PD conditions. White regions indicate non-significant oscillations (“n.s.”), i.e., those time-frequency regions for which the wavelet amplitudes are below the amplitude of a Poissonian process with the same mean. The black line represents the frequency with the maximum wavelet amplitude for each beta phase (smoothed with a moving average for display purposes). (F) Increase of gamma peak frequency and gamma spectral power with a higher mean firing rate of isolated GPe-TI. The increase of the firing was obtained by increasing the input to the isolated population. For comparison, the original nucleus spectral power and peak frequency are represented with a blue dashed line, for Parkinsonian condition.

We found that, under healthy conditions, the isolated GPe-TI firing spectrum closely resembled that of the full BG network within the gamma range (Figure 3B, top; R^2^=0.86 for frequencies between 50 and 150 Hz, see Methods). However, the isolated GPe-TI was unable to display beta oscillations, which are known to arise from the interplay among different BG nuclei ^58^. Under PD conditions, the behavior of the isolated GPe-TI diverged significantly from that of the full BG network (Figure 3B, bottom; R² = 0.34). As expected, the pathological beta hypersynchrony, which arises from interactions across BG nuclei, was entirely absent in the isolated GPe-TI. Furthermore, gamma activity in the isolated GPe-TI was notably diminished, showing reductions in both spectral power and peak frequency relative to the full BG network (Figure 3B). This suggests that, while self-inhibition was sufficient to sustain gamma oscillations under healthy conditions, additional inter-nuclei interactions are critical for maintaining both beta and gamma dynamics under pathological dopamine-depleted states.

To investigate this discrepancy further, we analyzed the effect of dopamine depletion on gamma power and peak frequency in both the isolated and full BG scenarios (Figure 3C). For low dopamine depletion levels (D_d_ < 0.87), the gamma power and peak frequency of the isolated and full BG GPe-TI populations remained similar, as indicated by a high goodness-of-fit between the firing spectra of the two scenarios (Figure S4A, left). However, beyond this threshold (D_d_ > 0.87), their behaviors began to diverge (Figure S4A, left): while gamma power and peak frequency increased in the full BG network, they progressively declined in the isolated GPe-TI. Interestingly, near this critical threshold (D_d_ ≈ 0.87), we observed that pathological beta power in the full BG network began to dominate and override gamma activity. This finding suggests that the observed differences between the isolated and full BG cases arise from the increasingly intense beta oscillations in the full network.

To support this hypothesis, we compared the GPe-TI firing rates across dopamine depletion levels (Figure S5). The average firing rates of the isolated and fully connected GPe-TI were comparable, decreasing progressively with dopamine depletion (Figure S5A, left). However, the standard deviation of the firing rates showed a marked divergence between the two scenarios (Figure S5A, right). In the isolated GPe-TI, the standard deviation decreased alongside the average firing rate, which led to a corresponding reduction in gamma power and peak frequency. This pattern reflected a direct relationship between firing dynamics and gamma oscillations in the absence of broader network interactions. In contrast, the full BG network exhibited a different behavior. While the average firing rate decreased with dopamine depletion, the presence of beta oscillations caused the standard deviation of GPe-TI firing rates to increase with higher levels of dopamine depletion (S5A Fig, right). These beta oscillations, driven by interactions across BG nuclei, induced significant fluctuations in GPe-TI firing rates, particularly at higher dopamine depletion levels (Figure 3D). Such fluctuations transiently elevated GPe-TI firing rates, leading to an increase in gamma power and peak frequency, an effect absent in the isolated GPe-TI scenario where beta oscillations were not present. To quantitatively explore the relationship between beta and gamma oscillations in the full BG model, we analyzed the beta-triggered scalogram of GPe-TI firing rates under PD conditions (Figure 3E). This analysis revealed that both gamma spectral power and peak frequency were strongly influenced by the phase of beta oscillations. Specifically, gamma activity reached its highest power and frequency at a beta phase of approximately 0, corresponding to the peak of GPe-TI firing observed in the full BG model (Figure 3D). Crucially, these fluctuations were absent in the firing of the isolated GPe-TI due to the absence of beta oscillations (Figure 3D and Figure S5A, right).

This coupling between gamma oscillations and beta phases explains the differences in gamma power and peak frequency observed in the PSDs of the isolated and full BG cases. The PSD provides an average power estimation over the entire time axis, disproportionately emphasizing the higher-frequency components of gamma oscillations, as these coincide with periods of elevated power driven by beta synchronization.

Consequently, the PSD of GPe-TI in the full BG model exhibited gamma oscillations of greater power and higher peak frequency compared to the isolated case (Figure 3B, C), driven by the transient beta-triggered increases in firing rates. To test whether the isolated GPe-TI could replicate these dynamics, we analyzed the relationship between its average firing rate and the resulting gamma power and frequency (Figure 3E). As expected, we found a positive correlation between these factors (firing rate and gamma peak frequency: Spearman ρ = 1.00, **p<0.001; firing rate and gamma power: Spearman ρ = 1.00, **p<0.001; gamma peak frequency and power: Spearman ρ = 1.00, **p<0.001), reinforcing the hypothesis that the enhanced gamma activity in the full BG model is attributable to transient increases of GPe-TI firing rates driven by beta oscillations.

Nonetheless, adjusting the average firing rate of the isolated GPe-TI revealed its limitations in fully replicating the spectral properties of the full BG model (Figure S4A, right). Even with an optimized GPe-TI firing rate that maximized the goodness-of-fit (maximum R²=0.80 for a firing rate of 45.22 spike/s), the match remained incomplete. This discrepancy highlights the critical role of beta-driven fluctuations and broader network interactions in shaping gamma oscillations (Figure S4A, right; maximum R2=0.80 for a firing rate of isolated GPe-TI equal to 45.22 spike/s).

Overall, these results underscore that GPe-TI self-inhibition is both necessary and sufficient for the emergence of gamma oscillations under both healthy and pathological conditions. However, in pathological conditions, beta-driven fluctuations (arising from network interactions beyond self-inhibition) play a critical role in modulating gamma activity, a dynamic that cannot be fully replicated by simply adjusting the nucleus’s average firing rate.

### D2 self-inhibition shapes gamma oscillations features

Following the same approach used for the GPe-TI nucleus, we systematically silenced each synaptic pathway individually in the full-BG model and analyzed its impact on D2 gamma power. Specifically, we calculated the ratio of D2 gamma power with each connection silenced to the gamma power in the intact full-BG model (Figure 4A). Since gamma oscillations in D2 neurons were absent in dopamine depletion levels corresponding to the healthy condition, this analysis was conducted under the PD condition. Our findings revealed that disrupting D2 self-inhibition nearly abolished gamma oscillatory activity (Figure 4A and Figure 4B). This disruption was accompanied by a significant increase in beta oscillation power (Figure 4B), suggesting that D2 self-inhibition plays a critical role in sustaining gamma oscillations in the PD state while simultaneously suppressing beta activity.

**Figure 4.**
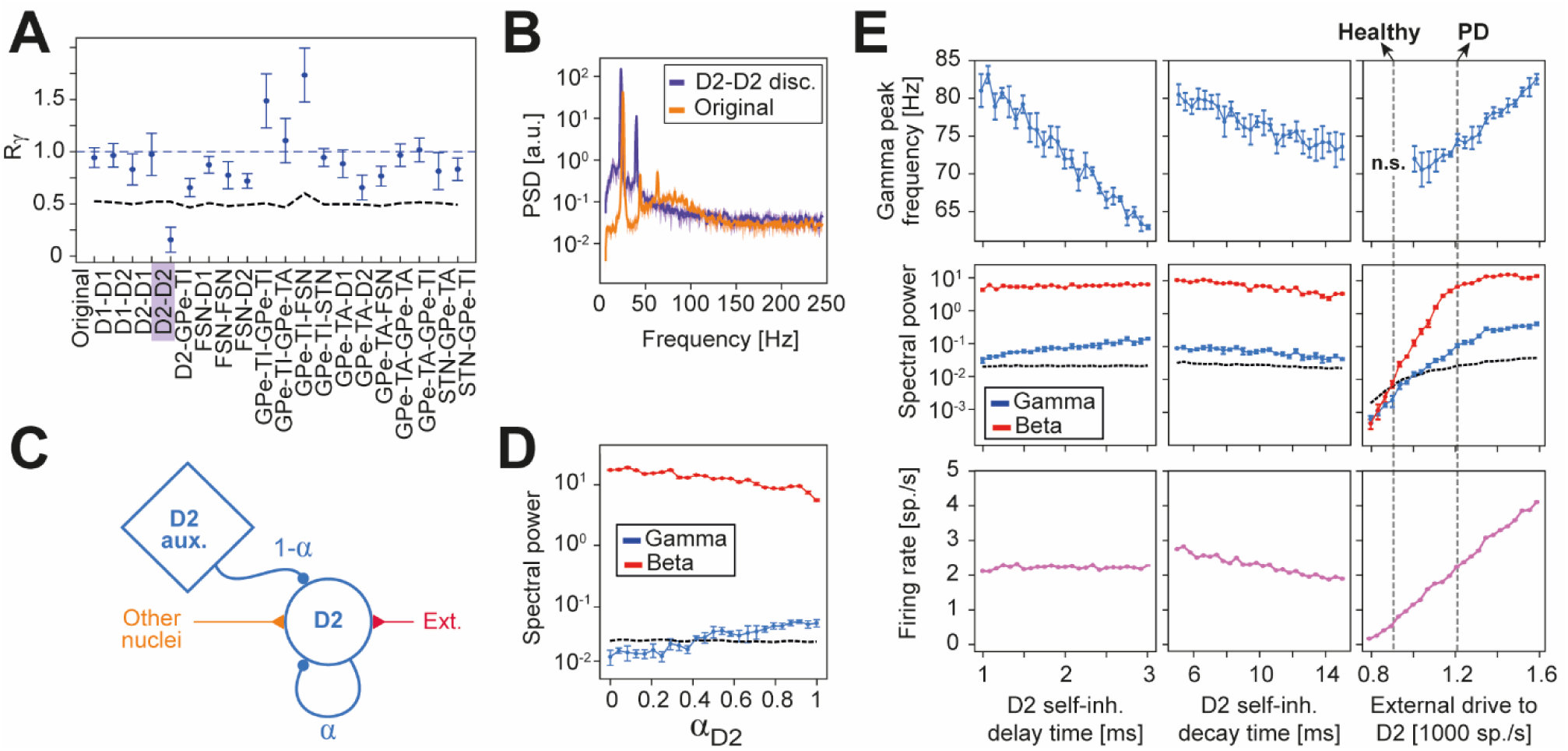
Self-inhibition parameters and external drive modulate D2 oscillations. (A) Ratio Rɣ of D2 gamma power with each synaptic connection silenced compared to the gamma power in the intact full-BG model. The spectral power ratio of D2 activity in the gamma band is shown with (‘original’) and without each synaptic connection in the full-BG model, with the specific connection removed indicated on the x-axis. For each removed connection, the threshold (black dashed line) for significant oscillations is computed as the ratio between the power of equivalent Poisson processes and the gamma power in the fully connected model. The blue dashed line indicates the spectral power ratio for the original fully connected case. (B) Power spectral densities (PSDs) of D2 with (orange) and without (purple) self-inhibitory connection in Parkinsonian condition (D_d_=1). (C) Scheme illustrating the procedure for reducing D2 self-inhibition: the weights of self-inhibitory connections were progressively reduced by a multiplicative factor α Є [0,1], and input from an auxiliary Poissonian population (D2 aux.) was added to compensate for the reduction in self-inhibitory input. The afferent activity from other BG nuclei and external inputs were left unchanged. (D) D2 spectral power of beta (red) and gamma (blue) bands across self-inhibition synaptic strength α. The threshold for statistically significant oscillatory power (dashed black line) was determined considering the power spectrum of an equivalent Poisson process with a matched firing rate. (E) Gamma peak frequency (top row), beta (red) and gamma (blue) spectral powers (middle row), and D2 firing rate (bottom row) across D2 self-inhibition delay (left column), D2 self-inhibition decay time (middle column), and the strength of external input to D2 (right column). In the middle row, the threshold for significant oscillation was indicated with a black dashed line. All data was reported as mean and standard deviation across five simulations. In the top row, peak frequencies are not reported for non-significant (“n.s”, according to comparison with the power of an equivalent Poissonian process with matched firing rate) gamma oscillations. The grey dashed lines show the value for which the external input match the healthy and pathological conditions.

To further investigate the role of D2 self-inhibition in regulating gamma oscillations, we systematically reduced the strength of self-inhibitory feedback and assessed its effect on D2 gamma power. To maintain a consistent average input to D2 despite the reduced self-inhibition, we introduced a compensatory Poisson input to offset the diminished inhibitory drive (see Methods and Figure 4C, where it is indicated with “D2 aux.”). Unlike what was observed for GPe-TI, this compensatory input was sufficient to maintain a stable D2 average firing across self-inhibition synaptic strengths (Figure S3B, left).

Our findings revealed that gradually reducing the strength of D2 self-inhibition (α_D2_) led to a progressive decrease in gamma oscillation power (Figure 4D). This decline continued until gamma oscillations completely disappeared when the D2-D2 synaptic strength was reduced to 40% of the original reference value (Figure 4D; Spearman ρ = 0.96, **p<0.001). At this point, gamma power fell below the threshold for statistical significance. A similar trend was observed for the D2 gamma peak frequency (Figure S3B, right; Spearman ρ = 0.80, **p<0.001). In contrast, beta oscillations in D2 neurons exhibited an opposing trend, with power increasing as the D2-D2 synaptic strength decreased (Figure 4B; Spearman ρ = −0.97, **p<0.001). This inverse relationship between gamma and beta oscillations in D2 neurons closely mirrored the results observed in GPe-TI.

Beyond synaptic strength, we investigated how other characteristics of D2 self-inhibition (specifically, synaptic decay time and delay) impacted gamma activity. Increasing the synaptic decay time of D2-D2 connections caused a slight decrease in gamma power (Figure 4E, middle; Spearman = −0.88, **p<0.001), gamma frequency (Figure 4E, top-middle; Spearman ρ = −0.95, **p<0.001), and the average firing rate of D2 neurons (Figure 4E, bottom-middle; Spearman ρ = −0.98, **p<0.001). This aligns with the notion that longer decay times reduce the overall excitability of the nucleus by prolonging inhibitory input, leading to a reduction in firing activity and, consequently, in gamma power and frequency.

In contrast, altering the synaptic delay of D2-D2 connections had minimal effect on the average firing rates of D2 neurons (Figure 4E, bottom-left; Spearman ρ = 0.11, p = 0.59). However, synaptic delay exerted distinct and opposing effects on gamma oscillations: gamma power increased (Figure 4E, middle-left; Spearman ρ = 0.98, **p<0.001), while gamma frequency decreased (Figure 4E, top-left; Spearman ρ = −0.99, **p<0.001). These findings suggest that synaptic delay primarily modulates the timing and efficiency of self-inhibitory interactions within D2 neurons, thereby affecting gamma oscillations more directly and independently of firing rate changes. This pattern mirrors the behavior observed in the GPe-TI, where changes in synaptic decay times affected firing rates and gamma oscillations in tandem, while changes in synaptic delay acted selectively on gamma features.

Finally, we assessed the impact of modulating external excitatory drive to D2 neurons on gamma oscillations, a manipulation that mimics the effects of dopamine depletion and models the transition from healthy to the Parkinsonian condition. As expected, increasing the excitatory drive to D2 neurons elevated their average firing rate (Figure 4E, bottom-right; Spearman ρ = 1.00, **p<0.001), which was accompanied by simultaneous increases in both gamma and beta power (Figure 4E, middle-right; gamma power: Spearman ρ = 1.00, **p<0.001; beta power: Spearman ρ = 0.96, **p<0.001) as well as in gamma peak frequency (Figure 4E, top-right; Spearman ρ = 0.90, **p<0.001). Interestingly, within the range of D2 external drive levels corresponding to healthy PD transition, both beta and gamma oscillations emerged as hallmarks of shifting network dynamics. Notably, beta oscillations appeared earlier during the transition from healthy to pathological state, while gamma oscillations became prominent only at later stages. These findings highlight the potential of striatal D2 gamma oscillation power and frequency as sensitive markers for tracking dopamine depletion and the progression between healthy and PD conditions.

### Gamma oscillations in D2 are modulated by beta activity

In the previous section, we demonstrated that D2 self-inhibition is a necessary condition for the emergence of gamma activity within the nucleus. Analogously to what was done for GPe-TI, we next investigated whether the D2 self-inhibitory loop alone was also sufficient for gamma rhythmogenesis. To address this, we analyzed an isolated D2 population configured with the same parameters as the original network but disconnected from all other basal ganglia nuclei. In this setup, inputs from other nuclei were replaced with auxiliary constant-rate Poisson inputs calibrated to match the firing activity of their corresponding original projections (Figure 5A, indicated with “FSN aux.”, “D1 aux.” and “GPe-TA aux.”).

**Figure 5.**
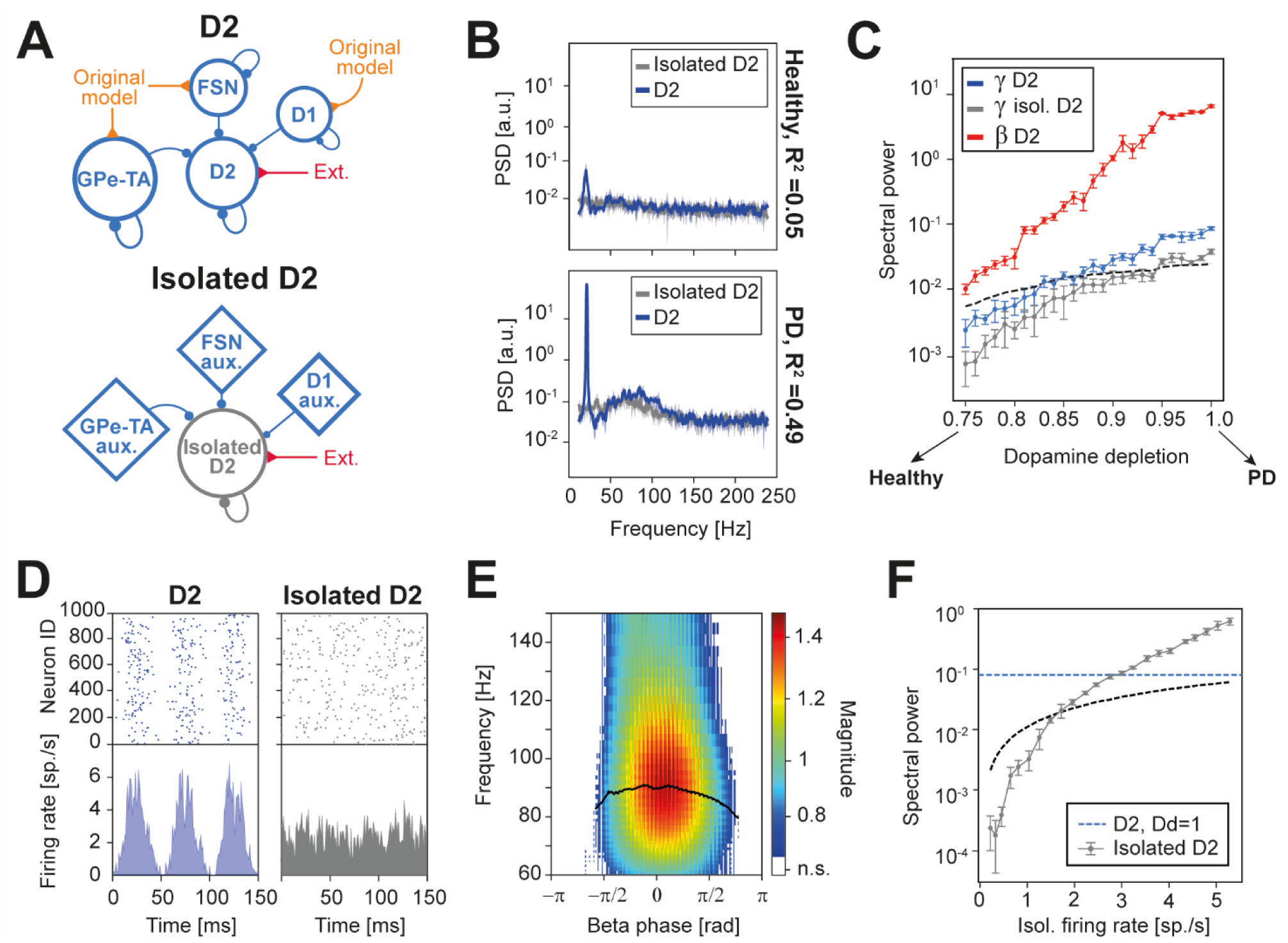
Interplay between beta and gamma oscillations in D2. (A) Schematic of isolation procedure for D2: the original nucleus (top) was detached by the rest of the model and the inputs from other nuclei were substituted with Poissonian sources (indicated with “aux.” in the bottom panel) with the same firing rates. External drive and self-inhibition were left unchanged. (B) Comparison of the power spectral densities of D2 (blue) and isolated D2 (grey) under healthy (D_d_=0.75, bottom) and Parkinsonian conditions (D_d_=1, bottom). (C) Spectral power in the gamma band of D2 (blue) and isolated D2 (grey) across D_d_. The spectral power in the beta band (red) and the threshold for significant oscillations (black dashed line), computed as the spectral power of a Poissonian process with the same firing rate, are also reported. Results are reported as mean across five simulations of 5 seconds. (D) Raster plots (top) and activity traces (bottom) of D2 (left) and isolated D2 (right), in parkinsonian condition. (E) Scalogram of D2 activity averaged across beta cycles under Parkinsonian conditions. White regions indicate non-significant oscillations (“n.s.”), i.e., those time-frequency regions for which the wavelet amplitudes are below the amplitude of a Poissonian process with the same mean. The black line represents the maximum wavelet amplitude frequency for each beta phase (smoothed with a moving average for display purposes). (F) Increase of gamma spectral power with a higher mean firing rate of isolated D2. The original nucleus spectral power for the Parkinsonian condition (blue dashed line) and the threshold for significant oscillations (black dashed line) are reported.

Under healthy conditions, we found that the spectral patterns of D2 firing activity in the isolated nucleus and the full BG network were poorly aligned (Figure 5B, R^2^=0.05), as neither scenario exhibited gamma activity. However, under PD conditions, gamma oscillations emerged in both the isolated D2 population and the full BG model. Despite this, the consistency between the two spectral patterns remained weak (Figure 5B, R^2^=0.49), with the isolated D2 population displaying reduced gamma power and central frequency compared to the full BG scenario. Furthermore, as expected, the pathological beta hypersynchrony observed in the full BG D2 was absent in the isolated case.

To further investigate this discrepancy, we analyzed the effect of dopamine depletion on gamma power in both the isolated D2 and full BG scenarios (Figure 5C). In the full BG model, increasing dopamine depletion led to a concurrent rise in both beta and gamma power. Notably, gamma oscillations in this scenario crossed the threshold of statistical significance at intermediate levels of dopamine depletion, approximately halfway through healthy and PD conditions. A similar dependence on dopamine depletion was observed in the isolated D2 nucleus, but the magnitude of gamma oscillations was consistently lower than in the full BG network. As a result, gamma power in the isolated D2 remained below the significance threshold until very high levels of dopamine depletion were reached. Accordingly, increasing dopamine depletion level led to only a mild improvement in the goodness-of-fit between the firing spectra of the scenarios (Figure S4B, left). To elucidate the mechanisms underlying the reduced gamma power in the isolated D2 nucleus, we explored the relationship between beta and gamma oscillations in the full BG model. This approach was motivated by the observation that beta oscillations played a key role in enhancing gamma activity in the GPe-TI (Figure 3). To quantitatively examine this relationship, we computed the beta-triggered scalogram of D2 firing rates under PD conditions (Figure 5E). This analysis was restricted to PD states because D2 gamma oscillations were absent in healthy conditions.

Our analysis revealed that both gamma spectral power and peak frequency were strongly modulated by the phase of beta oscillations. Specifically, gamma activity in the D2 population transitioned rhythmically between non-significant and significant power values as a function of the beta cycle phase. Consistent with our findings in the GPe-TI, D2 gamma power and frequency peaked at a beta phase corresponding to the maximum D2 firing rate observed in the full BG model (Figure 5D). In contrast, these beta-driven fluctuations were absent in the isolated D2 population due to the lack of beta oscillations (Figure 5D and Figure S5B, right).

These results indicate that the average firing rate of D2 neurons is insufficient to sustain gamma rhythmogenesis when considered in isolation. Instead, beta oscillations in the full BG model transiently elevate D2 firing rates, driving the nucleus into a firing regime where its self-inhibitory dynamics have sufficient energy to sustain gamma rhythmogenesis. This beta-mediated mechanism highlights the critical role of inter-nuclear interactions in modulating gamma activity under pathological conditions.

To determine whether the lack of gamma activity in the isolated scenario could be exclusively resolved by modulating D2 average firing activity, we analyzed the relationship between gamma power and the average firing rate of the isolated D2 population. As expected, increasing the average D2 firing rate led to a corresponding increase in gamma activity within the nucleus. Specifically, we found that an increase of 2.8 sp./s was required for the isolated D2 to match the gamma power observed in the full BG network. However, adjusting the firing rate of D2 alone was insufficient to fully replicate the spectral characteristics of gamma activity in the full BG circuitry (Figure S4B, right; maximum R^2^=0.70 for a firing rate of isolated D2 equal to 4.33 spike/s).

Overall, these findings demonstrate that D2 self-inhibition is necessary but not sufficient for gamma rhythmogenesis, under both healthy and pathological conditions. The emergence and modulation of gamma oscillations critically depend on beta-driven fluctuations, which arise from inter-nuclear interactions within the full basal ganglia circuitry. This underscores the importance of network-wide dynamics in shaping gamma activity and highlights the limitations of isolated nuclei in fully replicating the spectral properties of the full system.

## Discussion

### Nucleus-specific gamma rhythmogenesis in a basal ganglia spiking model

Using a computational spiking model of the basal ganglia ^58^, we identified two distinct types of gamma activity, originating from separate nuclei (Figure 1): D2 receptor-type medium spiny neurons in the striatum and the prototypical population of the globus pallidus. These two types of gamma oscillations exhibited different characteristics. D2 gamma oscillations occurred exclusively under pathological conditions, were confined to lower frequencies, and showed a marked increase in power with dopamine depletion. By contrast, GPe-TI gamma oscillations were observed consistently across healthy and pathological states, with their power remaining unaffected by dopamine depletion.

Despite these distinctions, our findings revealed a shared mechanistic requirement for both rhythms: their generation relies critically on recurrent inhibition within their respective nuclei, consistent with established theoretical accounts of gamma rhythmogenesis in inhibitory networks ^59^. The removal of inhibitory connections in the model completely abolished gamma oscillations in both cases (Figure 2 and Figure 4), underscoring the centrality of this mechanism. Furthermore, altering the synaptic time scales of self-inhibition modulated the spectral power and frequencies of these oscillations, further supporting the pivotal role of local inhibitory circuits in gamma activity patterns.

While recurrent inhibition proved necessary for gamma rhythmogenesis in both nuclei, it alone could not fully account for the observed gamma dynamics. In GPe-TI, self-inhibition was sufficient to generate gamma oscillations in the isolation experiment (Figure 3); however, in the full basal ganglia network the frequency and power of these oscillations were modulated by interactions with beta-frequency activity. In contrast, gamma rhythmogenesis in D2 MSNs required not only recurrent inhibition but also phase-amplitude coupling with beta oscillations, highlighting a more complex interplay of mechanisms in this population (Figure 5).

### Relationship to previous computational models of basal ganglia

The spiking network model presented in this work is largely based on a previously presented model by our group ^58^. In the previous work, we focused on the relationship between dopamine depletion and beta oscillations, hence gamma oscillations were not investigated. The version of the network presented in Ortone et al. ^58^, however, displayed high-frequency oscillations exceeding the range of the currently available experimental results (Table S2). The novel version of the network described in the present work accounts instead for both beta and gamma oscillations in basal ganglia. For a detailed comparison of the modifications introduced in the network see the dedicated 4.1.4 in the Methods and Table S3.

Our findings are consistent with prior computational models that identify the GPe as the primary source of basal ganglia gamma oscillations ^53–56^. These models similarly reported the limited dependence of GPe gamma oscillations on dopamine depletion levels, despite variations in the oscillation frequencies across studies. Among these, only Lindi et al.^54^ explicitly incorporated the two subpopulations of the GPe, aligning with our identification of the prototypical population as the driver of gamma activity.

The literature on striatal gamma oscillations presents a more heterogeneous picture. Findings from a model focusing exclusively on striatal populations ^57^ align with our results, identifying D2 MSNs as the primary source of gamma oscillations. In contrast, a different model ^53^ attributes gamma activity in the striatum to fast-spiking interneurons rather than D2 MSNs. This discrepancy may stem from fundamental differences in model design, particularly the absence of explicit modeling of D1 and D2 MSN subpopulations in the latter study. Such differences highlight the role of neuronal subtypes in determining complex dynamics of gamma oscillations in the striatum.

### Agreement with experimental findings on gamma oscillations

Experimental evidence from local field potential recordings in rats further supports our findings, with gamma oscillations observed in both the striatum ^43^ and in the GPe ^42–45,60^. GPe gamma oscillations have been suggested to arise locally ^45^ and striatal gamma oscillations have been found to correlate with beta power ^43^, consistently with our results.

Although calibrated using rat data, our model might provide insights into gamma oscillations observed in the STN of human patients ^34,35,41,61^. Under the hypothesis that LFPs in basal ganglia nuclei primarily reflect synaptic inputs, as proposed for cortical recordings ^62–64^, STN gamma activity can reflect an increase in GPe-TI firing (Figure 6). Specifically, our model suggests that: (i) increases in the mean firing rate of GPe-TI neurons lead to enhanced gamma-band power in their firing spectra (Figure 2, Figure 6A and 6B); and (ii) this gamma-patterned activity is subsequently transmitted to the STN, such that elevated GPe-TI firing rates are reflected as increased gamma power in the STN LFP (Figure 6C). This interpretation is consistent with experimental observations showing that gamma oscillations are enhanced during deep brain stimulation ^39^ and dopamine replacement therapy ^35,39^. In animal models of Parkinson’s disease, increased pallidal firing activity has been reported following dopamine injection in the basal ganglia ^65,66^, as well as following deep brain stimulation of the subthalamic nucleus ^67,68^. Our model provides a candidate mechanistic substrate for the emergence and amplification of gamma oscillations observed in downstream structures associated with this perturbation. Our model provides a candidate mechanistic substrate for the emergence and amplification of gamma oscillations observed in downstream structures associated with these perturbations. Our model provides a candidate mechanistic substrate for the emergence and amplification of gamma oscillations observed in downstream structures associated with this perturbation.

**Figure 6.**
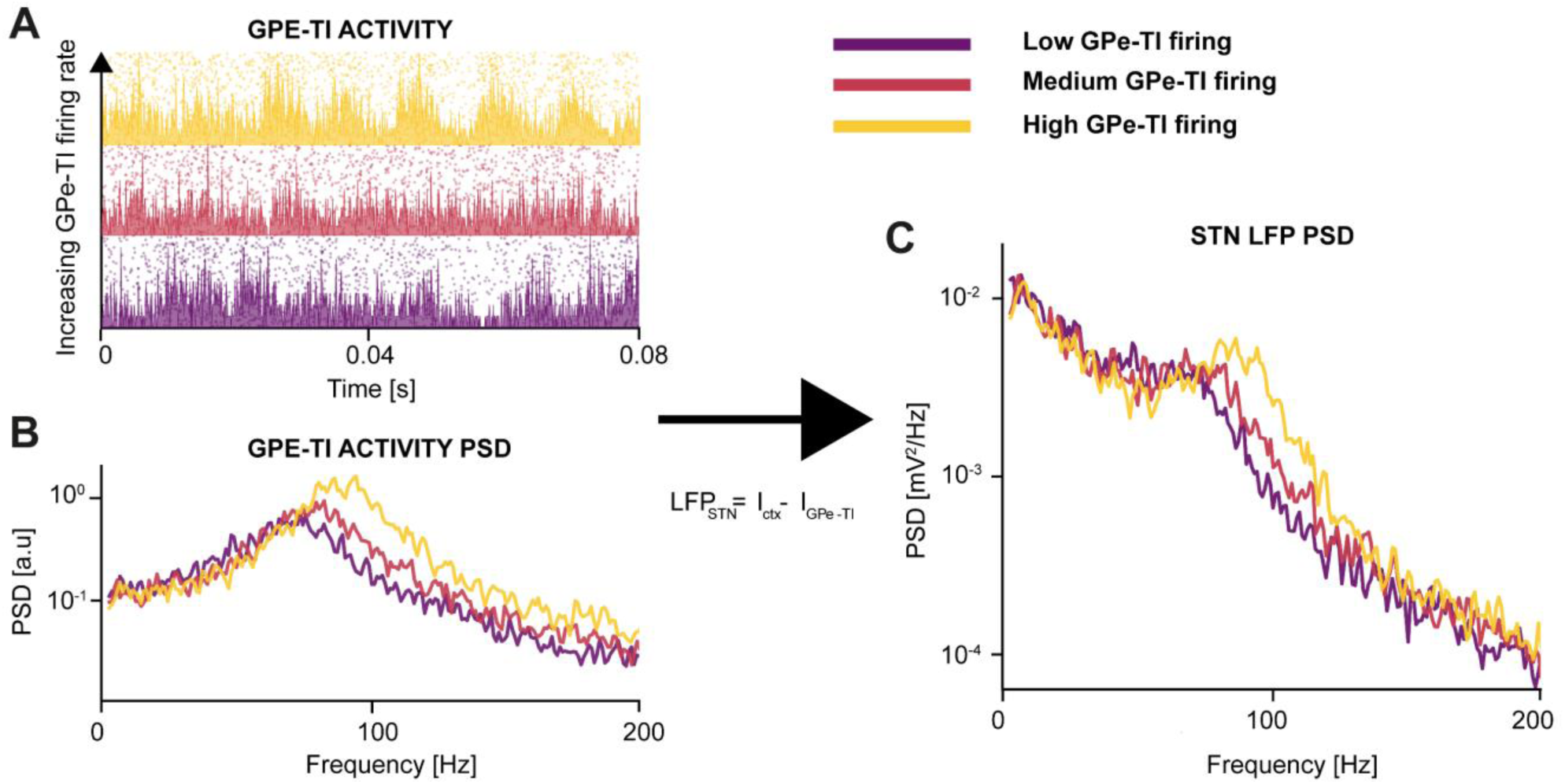
Pallidal firing rate drives gamma activity in subthalamic nucleus local field potential. (A) Firing activity of an isolated GPe-TI population for increasing mean firing rates [20–35 Hz]. (B) At higher mean firing rates, the power spectral density (PSD) of GPe-TI firing activity exhibits increasingly prominent gamma-band oscillations. (C) Assuming that the subthalamic nucleus (STN) local field potential (LFP) is generated as the sum of the absolute values of synaptic currents originating from GPe-TI and cortical inputs ^63^, gamma oscillatory activity emerges in the STN LFP, with power increasing as a function of the pallidal mean firing rate.

In contrast, striatal gamma oscillations are unlikely to directly affect STN local field potentials due to the lack of direct connections between these two nuclei ^2^.

A notable outcome of this study is the anti-correlation between pathological beta oscillations and GPe-TI self-inhibition. Interestingly, experimental recordings in 6-OHDA-lesioned rats have demonstrated an increase in GPe-TI self-inhibition under Parkinsonian conditions ^69^. This increase does not appear to result directly from changes in dopamine receptor activity within GPe-TI self-inhibition, suggesting that this enhanced synaptic strength reflects a form of adaptive homeostasis aimed at counterbalancing pathological beta oscillations.

The coupling between beta-phase and gamma-amplitude in the context of Parkinson’s Disease has been experimentally observed between the subthalamic nucleus and the motor cortex ^70^, within the motor cortex ^71,72^ and within the subthalamic nucleus ^70,73^, in both rodent models and human patients. Here, we characterized the mechanisms behind beta-gamma phase-amplitude coupling within the basal ganglia, demonstrating how beta-induced fluctuations of GPe-TI and D2 firing activities can influence gamma patterns. This provides a plausible explanation for the experimentally observed coupling between these two bands, shedding light on how beta oscillations can influence gamma-generating nuclei through intrinsic network interactions.

### Limitations of the study

Despite the strong alignment of our results with both experimental and computational studies, our computational framework has certain limitations. A key assumption in our study concerns the inputs provided to all populations within the BG circuitry, which were modeled as uncorrelated Poisson processes. While this approach simplified the complexity of network inputs, it does not account for the broader context in which the BG functions as part of an interconnected network involving the cortex and thalamus ^1^.

Evidence suggests that beta and gamma oscillations in the BG may be inherited from cortical activity ^74–76^. To achieve a more comprehensive understanding of these rhythms, future studies will then need to incorporate more biologically plausible models of cortical and thalamic inputs to capture their contributions to BG dynamics. Although such integrations were beyond the scope of this study, they represent an important direction for future work. Overall, our findings demonstrate that, within the assumptions of the present model, basal ganglia nuclei can intrinsically generate gamma oscillations, but does not exclude that extra-BG sources such as cortical or thalamic inputs play a role in shaping the basal ganglia activity within this band. Our modeling of dopamine depletion introduces another limitation. In this study, dopamine depletion was represented as an increased firing rate in the D2 populations, reflecting the heightened activity of the indirect pathway in PD ^50^. While effective in reproducing key features of pathological states, this simplification does not account for other critical factors, such as changes in synaptic strengths, plasticity mechanisms, or network-level adaptations that occur in Parkinson’s disease ^77^. By focusing on one direct consequence of dopamine depletion (namely, the imbalance between the direct and indirect pathways of the basal ganglia ^51^, see Figure S1) we captured qualitative changes in beta and gamma oscillatory activity, while acknowledging that additional dopamine-dependent mechanisms are required for a more complete biological representation. Future versions of the model could benefit from incorporating these additional factors to provide a more complete representation of the disease state.

Finally, lack of consensus in experimental literature remains in the estimation of precise values of the synaptic time scales, particularly those that have been shown to be critical for gamma-band modulation in the model. As illustrated in Figures 2 and 4, these parameters significantly affect both the power and the frequency of gamma oscillations. For this reason, we report in the manuscript a systematic parametric exploration of the synaptic decay time and synaptic delay for both connections. Through this analysis, we demonstrate that the model can reproduce a broad range of gamma oscillation frequencies depending on the selected synaptic time scales. Therefore, although uncertainty in the exact values of these parameters represents a relevant limitation of the present work, our results show that the model robustly captures the wide range of gamma peak frequencies reported in the experimental literature (Table S2), while maintaining consistent gamma oscillations across the explored parameter space.

### Conclusions

Our work identified two independent mechanisms supporting gamma oscillations in the basal ganglia. We found that these mechanisms are differentially affected by dopamine depletion and have different interactions with beta synchronization, highlighting a novel aspect of the pathological dynamics of basal ganglia in PD. Showing how pallidal gamma are reflected in STN LFP, we pave the way for a broader palette of neural markers of adaptive DBS including also the gamma band.

## Material and Methods

### Computational modeling

#### Description of the basal ganglia model

The basal ganglia model used in this work was adapted from a previous one implemented to study the origins of beta oscillations (Figure 1A and Ortone et al.^58^).

The model architecture comprised the striatum, the external globus pallidus and the subthalamic nucleus. The striatum was modeled with three neural populations: D1 receptor-type MSNs, D2 receptor-type MSNs and FSNs. The external globus pallidus was instead divided into two populations, according to experimental recordings in the pallidus of rats ^78,79^: the prototypic (GPe-TI) and arkypallidal populations (GPe-TA). The subthalamic nucleus was instead the only excitatory population of the model. The population numerosities of the model are reported in Table 1.

**Table 1.** Population numerosities of the six populations of the BG model.

| Population | FSN | D1 | D2 | GPe-TA | GPe-TI | STN |
| --- | --- | --- | --- | --- | --- | --- |
| N° of neurons | 420 | 6000 | 6000 | 264 | 780 | 408 |

The connectivities among populations were random with fixed probabilities based on previous works ^58,77^. Table 2 summarizes the connection properties within the BG model.

**Table 2.** Connection probabilities, synaptic delays and weights for all the connections within the BG model.

| Source | Target | Probability | Delay [ms] | Type | Synaptic weight [nS] |
| --- | --- | --- | --- | --- | --- |
| D1 | D1 | 0.0607 | 1.7 | Inh. | 0.12 |
| D1 | D2 | 0.0140 | 1.7 | Inh. | 0.30 |
| D2 | D1 | 0.0653 | 1.7 | Inh. | 0.36 |
| D2 | D2 | 0.0840 | 1.7 | Inh. | 0.25 |
| D2 | GPe-TI | 0.0833 | 7.0 | Inh. | 1.28 |
| FSN | D1 | 0.0381 | 1.7 | Inh. | 6.60 |
| FSN | FSN | 0.0238 | 1.0 | Inh. | 0.50 |
| FSN | D2 | 0.0262 | 1.7 | Inh. | 4.80 |
| GPe-TI | GPe-TI | 0.0321 | 1.8 | Inh. | 1.1 |
| GPe-TI | GPe-TA | 0.0321 | 1.0 | Inh. | 0.35 |
| GPe-TI | FSN | 0.0128 | 7.0 | Inh. | 1.60 |
| GPe-TI | STN | 0.0385 | 1.0 | Inh. | 0.08 |
| GPe-TA | D1 | 0.0379 | 7.0 | Inh. | 0.35 |
| GPe-TA | D2 | 0.0379 | 7.0 | Inh. | 0.61 |
| GPe-TA | FSN | 0.0379 | 7.0 | Inh. | 1.85 |
| GPe-TA | GPe-TA | 0.0189 | 1.0 | Inh. | 0.35 |
| GPe-TA | GPe-TI | 0.0189 | 1.0 | Inh. | 1.20 |
| STN | GPe-TA | 0.0735 | 2.0 | Exc. | 0.13 |
| STN | GPe-TI | 0.0735 | 2.0 | Exc. | 0.42 |

All populations in the model received external excitatory Poissonian inputs, simulating afferent drive from other brain structures. These inputs were calibrated to adjust each population’s firing rate within a range consistent with experimental observations (see Table S1). External input rates and synaptic strength are reported in Table 3.

**Table 3.** Parameters of the external excitatory input to each nucleus of the model. “*” indicates the value for healthy condition. For PD conditions, the excitatory input to D2 was set to 1200 spike/s (see “Modeling of dopamine depletion”).

| Target nucleus | $v_{ext}$ [spike/s] | Synaptic weight [nS] |
| --- | --- | --- |
| D1 | 1276 | 0.45 |
| D2 | 900* | 0.45 |
| FSN | 944 | 0.50 |
| GPe-TI | 600 | 0.25 |
| GPe-TA | 170 | 0.15 |
| STN | 500 | 0.25 |

Simulations were performed through the neural simulator ANNarchy ^80^. The integration method adopted was the 4th-order Runge-Kutta ^81^, with stepsize 0.1 ms. Each simulation of five seconds was repeated 5 times (except for the raster plots in Figure 3 and Figure 5, panel D). In all cases, we discarded an additional 1 second of simulation to avoid the effects of the initial conditions.

### Neuronal models

All the neurons in this work were modelled as adaptive conductance-based integrate-and-fire ^82,83^. Following the modeling approach described in Lindahl et al. and Ortone et al. ^58,77^, our model resulted from the combination of two previously implemented networks: the striatal model of M.D. Humphries et al.^84,85^ and the pallidus-subthalamic circuit model of Lindahl et al.^86^. Accordingly, it also inherited their choice of using two different point neuron models: quadratic integrate-and-fire for striatal populations and exponential integrate-and-fire for pallidus-subthalamic nuclei.

Specifically, striatal populations (D1, D2 and FSN) were modeled as adaptive quadratic integrate-and-fire ^87^:

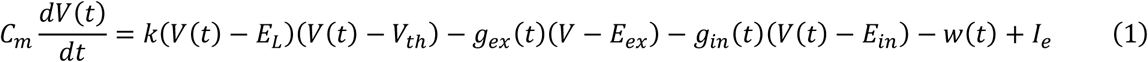

where k scales the voltage-dependent term, V is the membrane potential, E_L_ is the leak reversal potential, V_th_ is the threshold potential for spike initiation, g_ex_ and g_in_ are the excitatory and inhibitory conductances, E_ex_ and E_in_ are their respective reversal potentials, w represents the adaptation variable, and I_e_ is an external current.

Pallidal (GPe-TI and GPe-TA) and subthalamic neurons were considered instead as adaptive exponential integrate-and-fire ^87^:

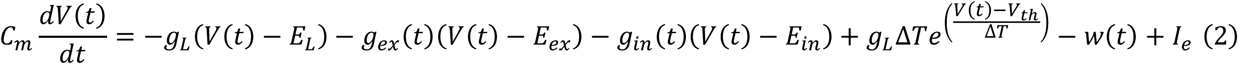

where C_m_ is the membrane capacitance, ΔT the slope factor determining the sharpness of the exponential activation near the threshold, and the exponential term represents the nonlinear exponential term modeling action potential generation.

For both models, if V>V_peak_ a spike is emitted and the membrane potential is reset to V_reset_.

In all cases but for FSN, the dynamic of the adaptive variable was described by:

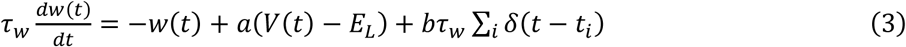

where *τ*_*w*_ is the adaptation time constant, *a* scales the subthreshold adaptation, *b* determines the strength of spike-triggered adaptation, and the sum accounts for discrete spike times of the neuron *t*_*i*_.

For FSN, according to the modeling of M.D. Humphries et al. ^84^, the dynamics for the adaptive variable was driven by:

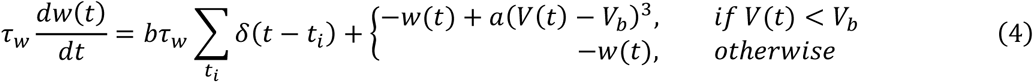

where the parameter V_b_ and the two regimes allow for a non-linear transition from silent to spiking dynamics.

The dynamics of excitatory and inhibitory synapses are described by the time evolution of conductances in each post-synaptic neuron, as represented by the following equations:

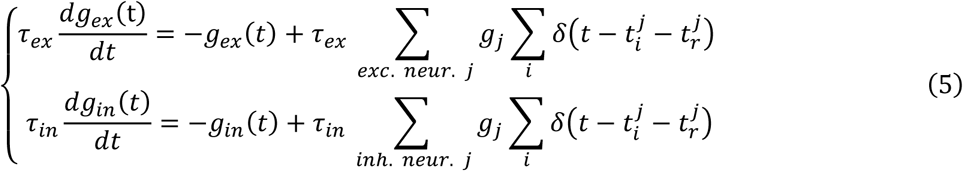

where *τ*_*ex*_ and *τ*_*in*_ represent the decay times for excitatory and inhibitory synaptic conductances of the postsynaptic neurons, respectively, and g_ex_ and g_in_ the corresponding synaptic conductances. g_j_ denotes the synaptic weight from the presynaptic neuron j, t^j^_i_ refers to the spike time i of the presynaptic neuron j, t_r_^j^ is the synaptic delay associated with the synapses originating from neuron j and *δ* is the Dirac delta function. The outer summations account for the contributions of all presynaptic neurons connected to the postsynaptic neuron under consideration, while the inner summations consider all spike events of the presynaptic neuron j.

All the neuronal parameters are equal to those of Ortone et al. ^58^, are reported in Table 4. The synaptic weights and delays can be found instead in Table 2.

**Table 4.** Parameters for the neuronal equation of each population of the BG model.

| Parameter | Unit | D1 | D2 | FSN | GPe-TI | GPe-TA | STN |
| --- | --- | --- | --- | --- | --- | --- | --- |
| $C_m$ | pF | 15.2 | 15.2 | 80.0 | 40.0 | 60.0 | 60.0 |
| $E_L$ | mV | -78.2 | -80.0 | -80.0 | -55.1 | -55.1 | -80.2 |
| $E_{ex}$ | mV | 0.0 | 0.0 | 0.0 | 0.0 | 0.0 | 0.0 |
| $E_{in}$ | mV | -74 | -74 | -74 | -65 | -65 | -84 |
| $\tau_{ex}$ | ms | 12.0 | 12.0 | 12.0 | 10.0 | 10.0 | 4.0 |
| $\tau_{in}$ | ms | 10.0 | 10.0 | 10.0 | 7.0 | 5.5 | 8.0 |
| $V_{th}$ | mV | -29.7 | -29.7 | -50.0 | -54.7 | -54.7 | -64.0 |
| $I_e$ | pA | 0.0 | 0.0 | 0.0 | 12.0 | 1.0 | 5.0 |
| $t_{ref}$ | ms | 0.0 | 0.0 | 0.0 | 0.0 | 0.0 | 0.0 |
| $V_{reset}$ | mV | -60 | -60 | -60 | -60 | -60 | -70 |
| $a$ | nS | -20 | -20 | 0.025 | 2.5 | 2.5 | 0.0 |
| $b$ | pA | 67.0 | 91.0 | 0.0 | 70.0 | 105.0 | 0.05 |
| $\tau_w$ | ms | 100.0 | 100.0 | 5.0 | 20.0 | 20.0 | 333.0 |
| $V_{peak}$ | mV | 40.0 | 40.0 | 25.0 | 15.0 | 15.0 | 15.0 |
| $\Delta T$ | mV | - | - | - | 1.7 | 2.55 | 16.2 |
| $g_L$ | nS | - | - | - | 1.0 | 1.0 | 10.0 |
| $k$ | nS/mV | 1.0 | 1.0 | 1.0 | - | - | - |
| $V_b$ | mV | - | - | -55 | - | - | - |

### Modeling of dopamine depletion

Our work aimed to investigate gamma oscillation patterns in healthy and Parkinsonian conditions. Parkinson’s disease has profound effects on striatal dynamics, which are particularly sensitive to the degeneration of dopaminergic neurons in the substantia nigra pars compacta ^88^. Experimental studies have shown that dopamine depletion alters the excitability of projection neurons in a receptor-specific manner ^89^. Specifically, neurons expressing D1 and D2 dopamine receptors exhibit opposite responses to dopaminergic inputs ^90,91^. In Parkinsonian conditions, this differential response results in an overall increase in the firing activity of D2 neurons, accompanied by a corresponding decrease in the firing activity of D1 neurons ^92,93^.

In our computational model, we directly implemented this increase in D2 neuron activity by modulating the rate of the external excitatory input, adopting the same assumption of Ortone et al.^58^. Specifically, the transition from healthy to PD was modeled with a dopamine depletion (D_d_) parameter according to the rule:

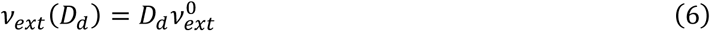

where the reference value 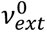 is equal to 1200 spike/s. The parameter D_d_ was then varied from 0.75 (representing healthy conditions) to 1 (for simulating the Parkinsonian condition).

In this way, the increase of dopamine depletion parameter D_d_ leads directly to an increase in D2 firing rate. Given the inhibitory nature of D2 neurons and their projections onto D1 neurons, enhancing D2 activity led to a corresponding reduction in D1 firing without the need for direct modulation of D1 external input (see Supplemental Text and Figure S1), effectively capturing the dynamics observed in PD without explicitly modeling dopamine signaling.

### Comparison with the model’s previous version

In this work, we used a newer version of the model compared to our previously developed one (53). The reason for this update was that, although the original model could generate gamma oscillations (an aspect not explicitly reported in Ortone et al. ^58^, see section **“**Relationship to previous computational models of basal ganglia” in Discussion), it exhibited excessively high frequencies of pallidal gamma activity, reaching approximately 130 Hz. While gamma oscillations were not the primary focus of the previous study, adjusting their frequency was essential for our investigation, as experimentally observed gamma activity in GPe-TI falls within a lower frequency range (Table S2). The revised parameter set effectively reduced the gamma oscillation frequency while preserving the beta rhythms and maintaining the original firing dynamics of the model.

The parameter adjustments primarily targeted the GPe-TI-GPe-TI and D2-D2 connections, as these pathways and their parameters significantly influence gamma oscillations (see sections “Globus pallidus self-inhibition drives gamma oscillations features” and “D2 self-inhibition shapes gamma oscillations features” in Results).

For GPe-TI self-inhibition the synaptic delay was set to 1.8 ms, and the decay time of all the connections to GPe-TI was increased from 5.5 ms to 7 ms. These changes reduced the central frequency of GPe-TI gamma oscillations by approximately 30 Hz while keeping the parameter values consistent with other computational models (Synaptic delay of GPe-TI to GPe-TI: ∼[2-4] ms ^94–96^; decay time of GPe-TI: ∼[5-8] ms ^96^). The synaptic weights for GPe-TI self-inhibition were reduced from 1.2 × 10⁻³ to 1.1 × 10⁻³, and the external input rate to GPe-TI neurons was decreased from 1530 spikes/s to 600 spikes/s. These adjustments were necessary to counterbalance the increase in gamma power caused by the lower synaptic delay (Figure 2E, left column).

For the D2-D2 connections, the synaptic weight was increased from 0.2 × 10⁻³ to 0.25 × 10⁻³, and the external input rate to D2 neurons was scaled by a factor of 1.11, enhancing the excitatory drive. Additionally, the external input to D1 neurons was increased from 1127 spikes/s to 1280 spikes/s. These changes were essential to restore the firing activity of D2 and D1 neurons and to recreate the gamma power of D2 gamma oscillations reported in Ortone et al. ^58^, which had been affected due to modifications in the GPe-TI pathway.

A summary of the changes made in our work as compared to Ortone et al. ^58^ is reported in Table S3.

### Analysis of the simulated signals

All spectral analyses were performed on the population firing rates, defined as the number of spikes produced by a population within a 1 ms time window divided by the product between the size of the population and the window length. Specifically, for the spectral analysis we computed the power spectral densities (PSDs) using Welch’s method ^97^, with 1 second of window length, 500 ms of overlap, and Hamming window.

If present, beta harmonics (see Figure 1B) were removed with a spectral interpolation technique ^98^, to avoid bias in the computation of gamma spectral power and frequency.

Although the power spectral densities from simulated data exhibit reduced complexity compared to experimentally measured ones, they still reflect spectral properties influenced by synaptic filtering, which introduced spurious trends unrelated to oscillatory dynamics ^99^. Since these trends could cause an overestimation in the computations of the spectral power, they were removed by fitting this aperiodic component and subtracting it. We chose to model the aperiodic component as a power law function of the frequency:

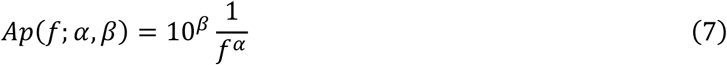

where *α* is the exponent of the power law and *β* an offset value. The fitting procedure was performed with the FOOOF method ^100^, which was used to identify the aperiodic components of the PSDs.

Once the beta harmonics and the aperiodic components were removed from the PSDs, we computed the oscillation frequency in both beta and gamma bands. This was estimated with the “centre of gravity” method ^101^:

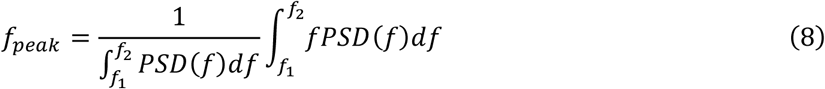

where we chose f_1_=10 Hz and f_2_=30 Hz for the beta band, f_1_=50 Hz and f_2_=150 Hz for GPe-TI gamma and f_1_=40 Hz and f_2_=120 Hz for D2 gamma. We adopted different ranges for the two gamma oscillations because GPe-TI and D2 gamma oscillations presented peaks at different frequencies (Figure 1B), and selecting specific ranges improved the accuracy of the peak identification.

We then used the computed central frequencies to choose the band on which to compute the mean spectral power:

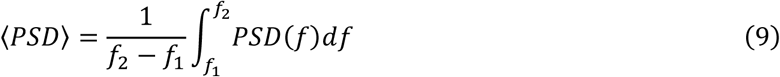

Specifically, we considered the beta and gamma bands respectively as 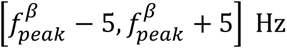 and 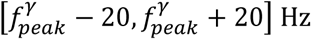. The use of bands centred around the peak frequencies was required to avoid changes in the spectral power due to the shift in frequency of the corresponding peak.

We also implemented a threshold to determine whether the power spectral densities displayed significant spectral power in the considered band. Specifically, the spectral power of the firing activity in the considered range was compared with that of fictitious Poissonian processes with the same average firing rate. To this purpose, 100 equivalent activities with a constant mean rate *v*(*t*) = *v*_0_ were simulated. We generated the number of spiking neurons n_j_ in each time bin j of the synthetic process from a binomial distribution:

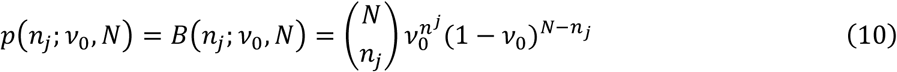

where N is the number of neurons in the considered population and *v*_0_ the probability of spiking each neuron, computed as the mean number of spikes of the population divided by the corresponding numerosity. We then estimated the threshold with the spectral power of the synthetic population in the chosen band, averaged across the 100 realisations of the process. If the mean spectral power of the original population exceeded this threshold, a significant oscillation was considered present in the activity. The thresholds were reported with dashed lines in all the figures representing analyses on the spectral powers.

### Computational experiments performed on the model

#### Identification of most relevant gamma oscillations

To identify which of the populations of the model presented significant gamma oscillations (section “Gamma oscillations in a computational model of basal ganglia” and Figure S1), we computed the gamma spectral power of activity of each nucleus, considering the gamma range as [50,120] Hz. Unlike all the others, for this analysis we adopted a fixed range for the gamma band to avoid any assumption about potential gamma oscillations. Furthermore, we selected a broad range of frequencies to capture any source of gamma activity and to account for the wide distribution of the observed gamma peaks. We then compared the spectral powers with those of the Poissonian processes in the same band and considered significant oscillations only those that exceeded this threshold (see Materials and Methods). With this approach, we found that two populations displayed gamma oscillations in their activities: GPe-TI and D2 (See Figure 1B and Figure 1C). Consequently, all the other analyses of this work were performed exclusively on these two populations.

##### 4.3.2. Identification of necessary projections for gamma oscillations

We aimed to determine whether gamma rhythms in the BG model depended on specific synaptic connections. To investigate this, we systematically disconnected each projection in the model and analyzed how gamma oscillations in the two nuclei were affected (Figure 2B and Figure 4B). Synaptic disconnection was implemented by setting the corresponding synaptic weights to zero. However, this removal altered the input received by the target population, potentially affecting not only the spectral pattern of its firing dynamics but also its average level. To compensate for the missing input while isolating the role of the connection in generating gamma oscillations, we introduced an auxiliary Poissonian population. This population matched the size and average firing rate of the original source population and provided a white noise input to the target nucleus through a projection with the same characteristics as the removed one. For instance, if the disconnected projection was from D2 to GPe-TI, an auxiliary Poissonian population with the same size and average firing rate as D2 was created to send input to GPe-TI through a projection with synaptic properties identical to the original "D2 to GPe-TI" connection. By maintaining a constant average input to the target nucleus, this approach ensured that overall firing dynamics remain stable within the BG model. However, it effectively eliminated any temporal structure originally present due to the disconnected synaptic projections. This allowed us to determine whether the specific temporal patterns of firing activity originating from a given projection contributed to the generation of gamma oscillations or whether their influence was merely due to the average input level of the source nucleus.

To compare the spectral power in GPe-TI (Figure 2B) and D2 (Figure 4B) with and without each connection, we then calculated the ratio R*γ* of gamma power with each connection silenced to the gamma power in the intact model. In this way, we were able to identify those projections whose presence was the most relevant to gamma rhythmogenesis.

### Investigation of the effects of recurrent synaptic parameters on oscillations

The disconnection of the model projections described in the previous subsection allowed for the identification of self-inhibition as necessary condition for gamma oscillations (Figure 2B and Figure 4B). To further characterize the role of these connections (i.e., D2 to D2 and GPe-TI to GPe-TI), we investigated the effects of their synaptic parameters on gamma oscillations (sections “Globus pallidus self-inhibition drives gamma oscillations features” and “D2 self-inhibition shapes gamma oscillations features” in Results). First, we analyzed the effects of reducing self-inhibitory strength in D2 and GPe-TI on the spectral power of both beta and gamma oscillations. This was achieved by scaling down the synaptic weights of recurrent inhibitory connections by a multiplicative factor α ∈ [0,1]. Since this reduction weakened self-inhibition and consequently altered the firing dynamics of the nucleus, we introduced an auxiliary Poissonian population (labeled ‘aux.’ in Figure 2C and Figure 4C) to compensate for the loss of inhibitory input. This auxiliary population matched the original nucleus in terms of neuron count, synaptic parameters, and average firing rate. It provided external input through a projection with synaptic weights scaled by 1−α relative to the original self-inhibition, ensuring that the overall level of inhibitory input remained unchanged. However, because the auxiliary population generated Poisson-distributed activity, this substitution eliminated the temporal structure inherent to the original self-inhibitory projections. This approach allowed us to isolate the role of recurrent inhibition in shaping gamma oscillations while preserving the nucleus’s mean input.

Next, we examined the effects of altering the synaptic delay and decay time of self-inhibition in GPe-TI and D2 on their firing rates, gamma spectral power, and gamma peak frequencies (Figure 2E and Figure 4E, left and center columns). We also studied how the change in the average rate of the external excitatory drive to GPe-TI and D2 affected these quantities (Figure 2E and Figure 4E, right column).

All relationships between the alteration of synaptic parameters and the observed quantities were evaluated using the Spearman coefficient, with the corresponding p-value computed via a bootstrap permutation test with 1,000 permutations. The investigations were performed in the pathological case (D_d_=1), because in healthy conditions only GPe-TI gamma oscillations were present.

### Isolation of gamma-generating nuclei

In addition to investigating whether self-inhibitory connections were necessary for gamma oscillations, we examined whether they were also sufficient to sustain gamma activity in GPe-TI and D2 (see sections “Gamma oscillations in the globus pallidus are modulated by beta activity” and “Gamma oscillations in D2 are modulated by beta activity” in Results). To this end, we created isolated versions of GPe-TI and D2, generating populations (“isolated GPe-TI” and “isolated D2” in Figure 3A and Figure 5A, respectively) that retained the same parameters and neuron count as in the full model. These populations were disconnected from all other nuclei while preserving their recurrent inhibitory connections and external excitatory Poissonian drive.

A key challenge of this isolation was to maintain firing rates comparable to those in the full model despite these removed inputs. To achieve this, we replaced the disconnected projections with inputs from auxiliary Poissonian populations. Similarly to what was detailed in previous sections, these auxiliary populations generated Poisson-distributed synaptic inputs with the same average firing rate and synaptic properties as those of the original source populations they replaced. This approach ensured that the isolated nuclei received the same average input as in the full model while eliminating any temporal (and thus spectral) structure present in the original projections. For instance, in the full BG model, GPe-TI receives inputs from GPe-TA, D2, and STN. In the isolated GPe-TI condition, these inputs were replaced by Poissonian versions of these nuclei, labeled GPe-TA aux., D2 aux., and STN aux. (Figure 3A). This substitution preserved the average level of synaptic inputs while preventing any spectral influence present in the original network activity, allowing us to assess whether self-inhibition alone was sufficient to generate gamma oscillations. We then compared the spectra of the isolated populations with the corresponding original ones, in both healthy and PD (Figure 3B and Figure 5B). Specifically, we computed the coefficient of determination R^2^ between the two power spectral densities (original and isolated), considering only the frequencies between 50 Hz and 150 Hz to avoid bias induced by the presence of beta oscillations. The PSDs considered for this analysis were averaged across five simulations, to limit noise variability.

The isolated populations were also used to compute the relationship between the firing rate of the population and the spectral power and peak frequency of gamma oscillations (Figure 3F and Figure 5F).

### Beta-induced modulations on gamma oscillations

Analysis of the population activity in nuclei exhibiting gamma oscillations suggested a potential coupling between beta and gamma rhythms (Figure 3D and Figure 5D). To explore this interaction, we conducted a time-frequency decomposition of the firing activities by calculating their beta-averaged scalogram (Figure 3E and 5E). Specifically, we first applied a Morlet wavelet transform (5 cycles, 200 frequency steps between 60 and 150 Hz) to the firing activity of each nucleus to obtain its time-resolved spectral power. Next, we extracted the beta phase by applying a Hilbert transform to the bandpass-filtered signal (Butterworth filter, 5th order, [10–30] Hz). The beta phase was then discretized into 100 bins, and the scalogram was averaged within each phase bin. This procedure allowed us to quantify how the power of different frequencies varied across the beta cycle, providing insight into the phase-dependent modulation of gamma oscillations.

To distinguish genuine gamma oscillations from background activity, we compared these amplitudes to those obtained from a Poisson process with the same average firing rate as the original nucleus. Specifically, we computed the wavelet transform of the firing rate of a Poissonian spiking process with the same duration and mean rate as the original data. The resulting gamma-band wavelet amplitudes were then used as a significance threshold: only frequency-phase bins where the original amplitude exceeded the Poisson-derived values were considered indicative of significant oscillatory activity.

## Acknowledgments

F.F. and M.A. were supported by the Italian Ministry of University and Research, under the complementary actions to the NRRP “Fit4MedRob—Fit for Medical Robotics” Grant (# PNC0000007). N.M. was supported by #NEXTGENERATIONEU (NGEU) and funded by the Ministry of University and Research (MUR), National Recovery and Resilience Plan (NRRP), project THE (IECS00000017) - Tuscany Health Ecosystem (DN. 1553 11.10.2022)". The work was also funded by “Fondo di beneficenza ed opere di carattere sociale e culturale” granted by “Banca Intesa San Paolo” in the context of the project ONDA (“Origini neurali delle disfunzioni dell’arto superiore su modello murino di parkinson”), CUP B53C22007580007. The work was also supported by #NEXTGENERATIONEU (NGEU) and funded by the Ministry of University and Research (MUR), National Recovery and Resilience Plan (NRRP), project EBRAINSItaly (IR0000011) - European Brain ReseArch INfrastructureS-Italy (DN. 101 16.06.2022).

## Author contributions

**F.F.:** Conceptualization, Formal Analysis, Software, Visualization, Writing – Original Draft Preparation; **M.A.:** Conceptualization, Software, Supervision, Writing – Original Draft Preparation; **N.M.:** Conceptualization, Supervision, Visualization, Writing – Original Draft Preparation; **E.C.:** Conceptualization, Project Administration, Supervision, Writing – Review & Editing; **A.M.:** Conceptualization, Funding Acquisition, Project Administration, Supervision, Writing – Original Draft Preparation, Writing – Review & Editing.

## Data and code availability statement

The codes necessary to reproduce the data that support the findings of this study are openly available at https://github.com/CNELab/Gamma-Oscillations-Basal-Ganglia.

## Declaration of generative AI and AI-assisted technologies in the manuscript preparation process

During the preparation of this manuscript, the authors used ChatGPT (OpenAI) to assist with language refinement and improvement of readability. All AI-assisted output was critically reviewed and edited by the authors, who take full responsibility for the content of the published article.

## Supplementary Material

**Table S1.** Experimental literature data on basal ganglia nuclei firing rates and corresponding model values under healthy conditions.

| Nucleus | Experimental firing rate | References | Model firing rate [sp./s] |
| --- | --- | --- | --- |
| D1 | 0.5-3 | 102,103 | $2.41 \pm 0.01$ |
| D2 | 0.5-3 | 102,103 | $0.526 \pm 0.004$ |
| FSN | 10-20 | 104,105 | $17.66 \pm 0.05$ |
| GPe-TI | 30-60 | 79,106 | $45.58 \pm 0.03$ |
| GPe-TA | 10-20 | 106 | $15.46 \pm 0.03$ |
| STN | 10-20 | 107,108 | $19.44 \pm 0.01$ |

**Table S2.** Frequency of gamma oscillations with the basal ganglia in experimental literature.

| Gamma oscillations central frequency | Source nucleus | Humans / rats | Reference |
| --- | --- | --- | --- |
| ~100 Hz | Striatum | Rats | 42 |
| ~70 Hz | Striatum | Rats | 60 |
| 80-100 Hz | Striatum | Rats | 43 |
| 80-110 Hz | Striatum and External | Rats | 44 |
| 80-110 Hz | Subthalamic Nucleus | Rats | 42 |
| 60-90 Hz | Globus Pallidus | Rats | 45 |
| 60-90 Hz | Subthalamic Nucleus | Human | 48 |
| 60-90 Hz | Subthalamic Nucleus | Human | 35 |
| 60-90 Hz | Subthalamic Nucleus | Human | 40 |

**Table S3.** Modified parameters in the updated model.

| Parameter | Previous model value | New model value |
| --- | --- | --- |
| GPe-TI to GPe-TI synaptic delay | 1 ms | 1.8 ms |
| GPe-TI synaptic decay time | 5.5 ms | 7 ms |
| GPe-TI to GPe-TI synaptic weight | $1.2 \times 10^{-3}$ | $1.1 \times 10^{-3}$ |
| GPe-TI external input rate | 1530 spikes/s | 600 spikes/s |
| D2 to D2 synaptic weight | $0.2 \times 10^{-3}$ | $0.25 \times 10^{-3}$ |
| D2 external input rate in PD conditions | 1083 spikes/s | 1200 spikes/s |
| D1 external input rate | 1127 spike/s | 1280 spikes/s |

### Supplemental Text

To demonstrate that modulating the external input to D2 is equivalent to jointly modulating the external inputs to D1 and D2, we examined the firing rates of D1 and D2 neurons across a two-dimensional range of external inputs to both populations (Figure S1A–B). In the model, the healthy and parkinsonian conditions are defined by characteristic firing rates: in the healthy state, D1 neurons fire at 2.5 spikes/s and D2 neurons at 0.5 spikes/s, whereas in the pathological state these values are inverted. We identified the combinations of external inputs to D1 and D2 that reproduce these firing-rate pairs, yielding four corresponding isocurves, and selected their intersections as representative points of the healthy and pathological conditions.

We then parameterized the line connecting these two points, which defines a continuous transition between the healthy and pathological regimes in the two-dimensional input space. This line can be expressed using a single scalar parameter t ∈ [0,1], as:

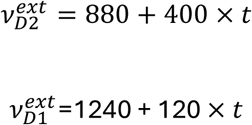

To verify that this single parameterization (corresponding to the simultaneous modulation of D1 and D2 inputs) produces network dynamics comparable to those obtained using the original parameter *D*_*d*_, we show in Supplementary Figure S1C–H that the firing rates of all six neuronal populations are consistent across the two parameterizations. This demonstrates that the original two-dimensional modulation of D1 and D2 external inputs can be reduced to a one-dimensional trajectory without significant loss of information about the firing dynamics of the model.

**Figure S1.**
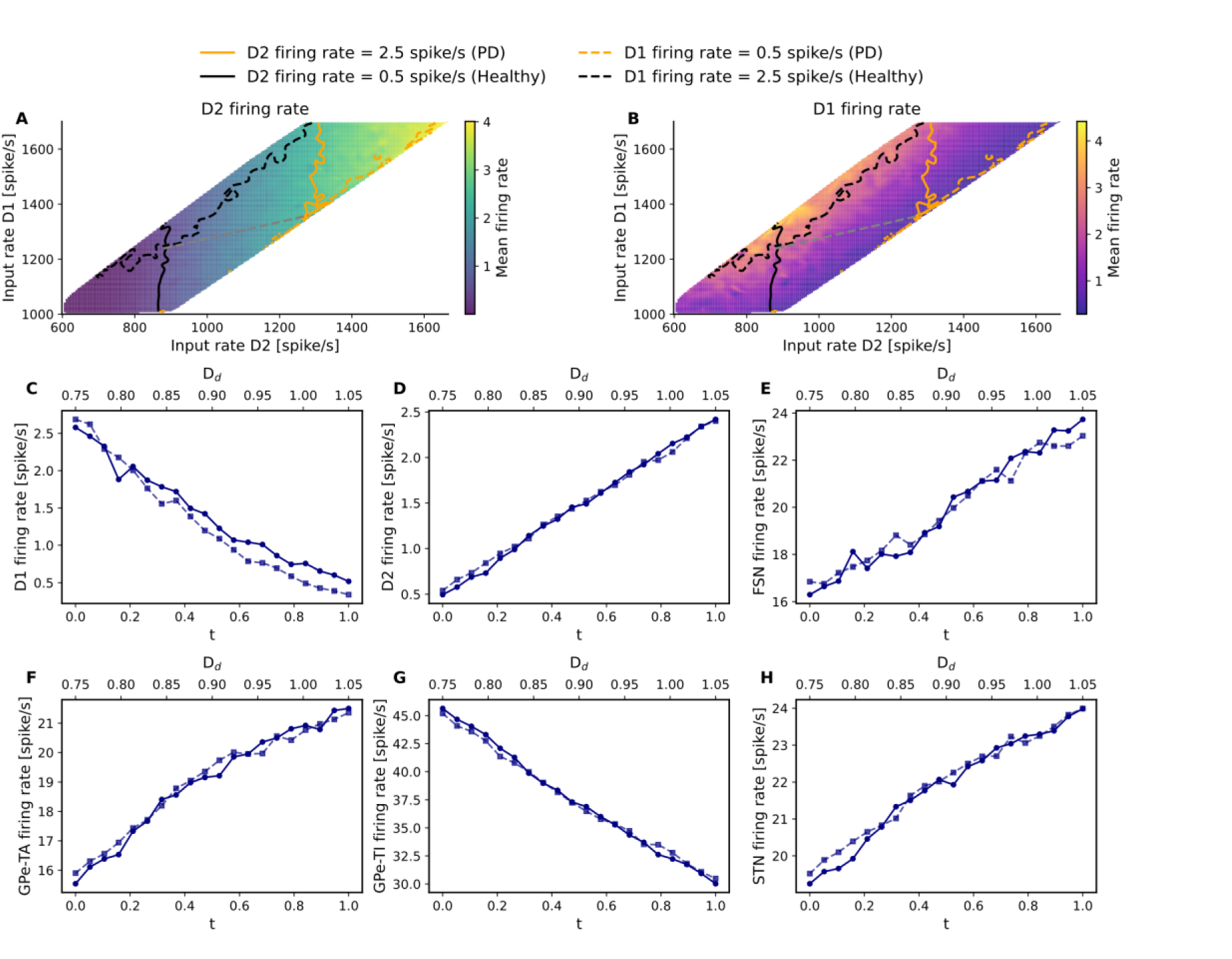
Mean firing rates of D2 (A) and D1 populations (B) as a function of the rate of the external input provided to D1 and D2 neurons. The black contours indicate healthy firing-rate levels of 2.5 spikes/s for the D1 population and 0.5 spikes/s for the D2 population. The orange contours indicate pathological (PD) firing-rate levels of 2.5 spikes/s for the D2 population and 0.5 spikes/s for the D1 population, respectively.. The grey dotted line denotes the range of firing rates explored by the model when dopamine depletion is modulated exclusively through the D_d_ parameter. Panels C–H show the firing rates explored by the parametrization of the D_d_ parameter (solid lines with circular markers) and by the reparameterization using t as a common parameter modifying, respectively, for the D1, D2, FSN, GPe-TA, GPe-TI, and STN populations.

**Figure S2.**
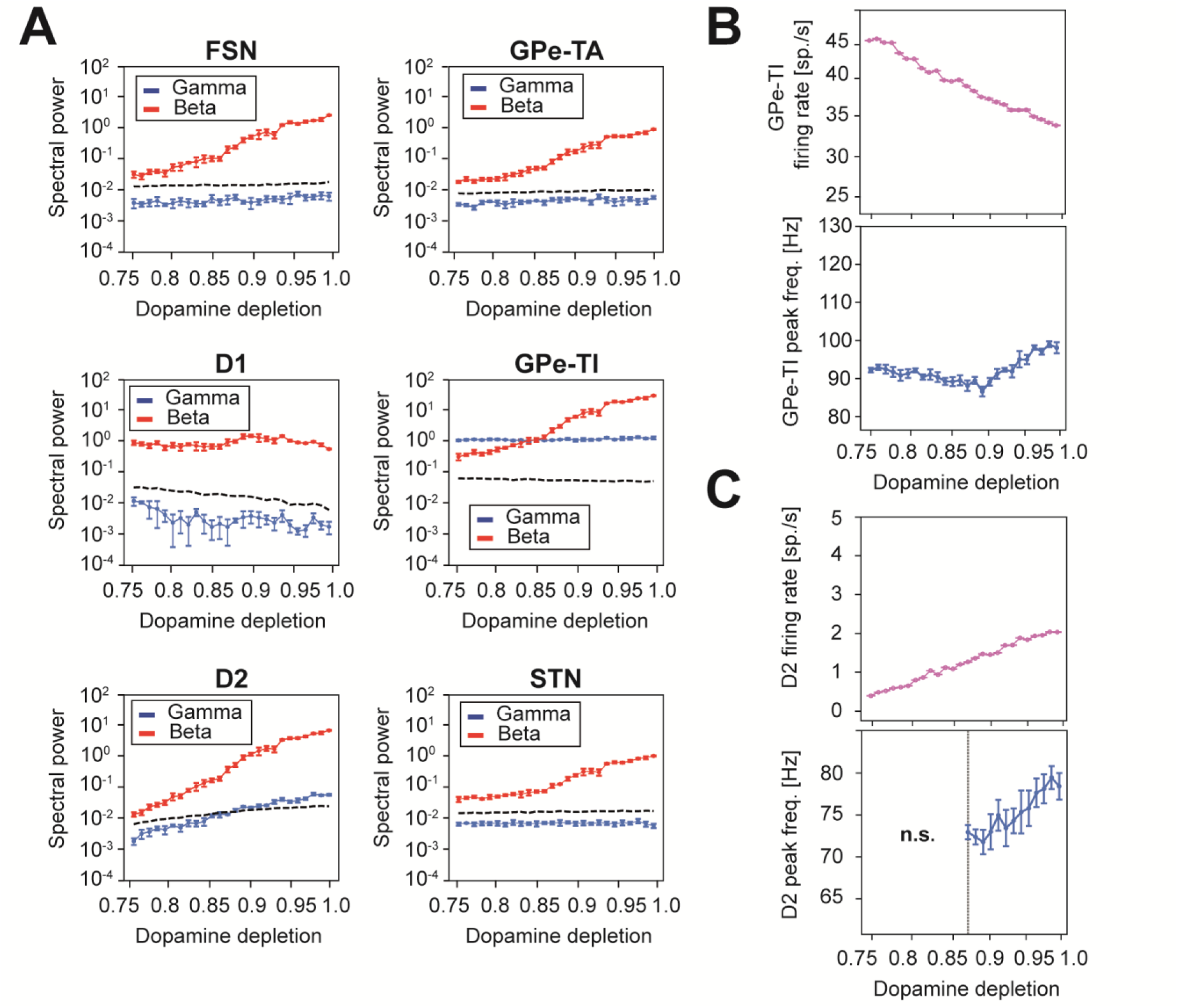
GPe-TI and D2 are the only nuclei displaying gamma activity. A) Spectral power in the gamma (blue) and beta (red) bands for each basal ganglia nucleus of the model, across different dopamine depletion levels. The threshold for significant gamma oscillations (black dashed line) is computed as the gamma spectral power of a Poissonian process with the same firing rate. Results are presented as mean and standard deviation across five simulations of 5 seconds. B) GPe-TI firing rate (top) and gamma peak frequency (bottom) with different levels of dopamine depletions. C) D2 firing rate (top) and gamma peak frequency (bottom) with different levels of dopamine depletions. Peak frequencies are not reported for non-significant (“n.s”, according to comparison with the power of an equivalent Poissonian process with matched firing rate) gamma oscillations.

**Figure S3.**
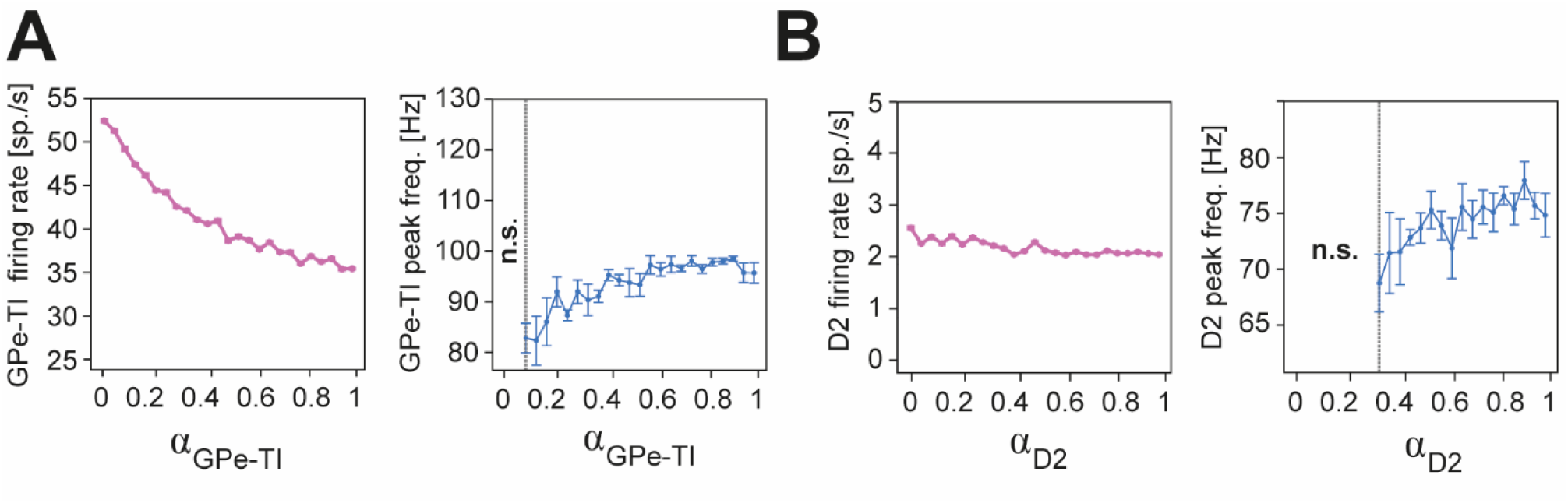
Effects of self-inhibition disconnection on firing rates and gamma peak frequencies of GPe-TI and D2. A) GPe-TI gamma peak frequency (left) and firing rate (right) across different levels of self-inhibition synaptic strength α_GPe-TI_. Peak frequencies are not reported for non-significant (“n.s”, according to comparison with the power of an equivalent Poissonian process with matched firing rate) gamma oscillations. Results are presented as mean and standard deviation across five simulations of 5 seconds. B) D2 gamma peak frequency (left) and firing rate (right) across different levels of self-inhibition strength α_D2_. Peak frequencies are not reported for non-significant (“n.s”, according to comparison with the power of an equivalent Poissonian process with matched firing rate) gamma oscillations. Results are presented as mean and standard deviation across five simulations of 5 seconds.

**Figure S4.**
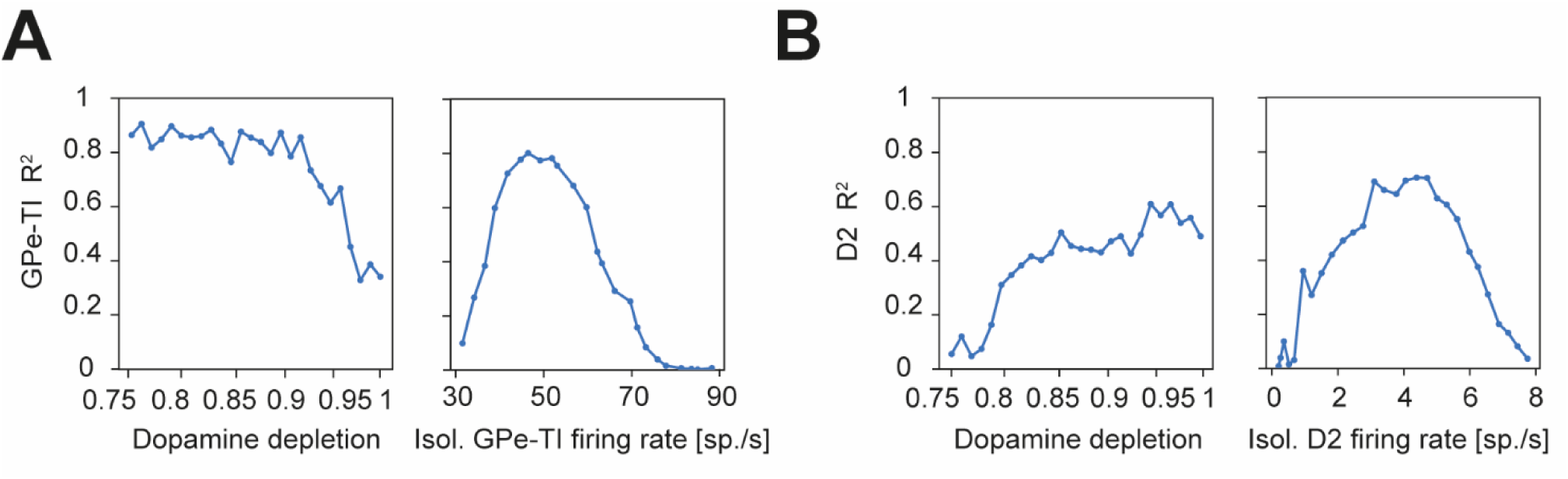
Similarity between the isolated and original power spectral densities across dopamine depletion levels and firing rates of the isolated nuclei. A) left, coefficient of determination R^2^ between the power spectral densities (considering the range [50-150] Hz) of the isolated GPe-TI and the full BG GPe-TI across dopamine depletion levels; right, R^2^ between the power spectral densities of GPe-TI at D_d_=1 and isolated GPe-TI, varying the firing rate of the latter and considering the range [50-150] Hz. B) left, R^2^ between the power spectral densities (considering the range [50-150] Hz) of D2 and isolated D2, across different levels of dopamine depletion; right, R^2^ between the power spectral densities of D2 at D_d_=1 and isolated D2, varying the firing rate of the latter and considering the range [50-150] Hz.

**Figure S5.**
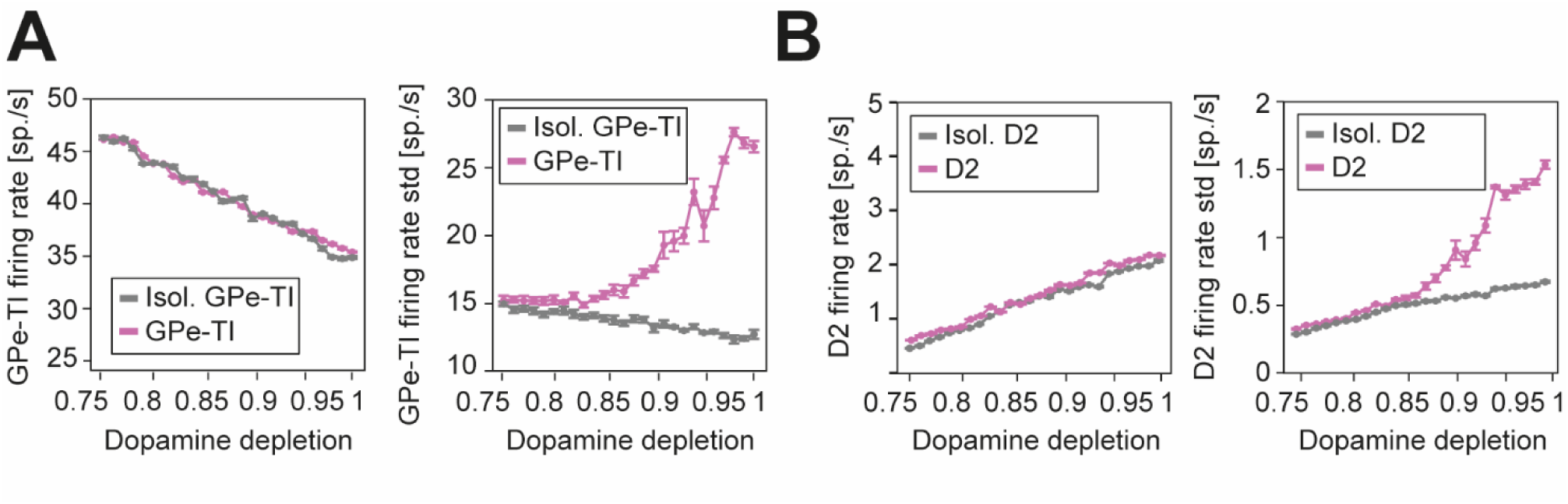
Isolated versions of D2 and GPe-TI present the same average firing activities of the original nuclei, but smaller fluctuations. A) Mean firing rate (left) and its simulation standard deviation (right) of GPe-TI (pink) and isolated GPe-TI (grey) across dopamine depletion levels (D_d_). The isolated GPe-TI population represents an equivalent of GPe-TI, but with inputs from other nuclei replaced by Poissonian sources with matching firing rates. Results are presented as the mean and standard deviation across five simulations, each lasting 5 seconds. B) Mean firing rate (left) and its simulation standard deviation (right) of D2 (pink) and isolated D2 (grey) across dopamine depletion levels (D_d_). The isolated D2 population represents an equivalent of D2, but with inputs from other nuclei replaced by Poissonian sources with matching firing rates. Results are presented as the mean and standard deviation across five simulations, each lasting 5 seconds.

